# A flexible Bayesian method for estimating stratigraphic intervals and their co-occurrence in time

**DOI:** 10.1101/2025.02.13.638199

**Authors:** Gustavo A. Ballen

## Abstract

The fossil record is both incomplete and subject to a number of biases with different sources (e.g., preservational, statistical), which hinders making inferences about the past of life on earth. Stratigraphic intervals allow estimate two important quantities from the fossil record of a given lineage, its origination, and extinction times. Many models are available in the literature, although specialising in implementation and therefore resulting in limitations of application. This toolset is extended herein by proposing a flexible method which allows one to estimate origination and extinction times, as well as model preservation potential in flexible ways. The present method is more general than previous alternatives, many of which constitute special cases. A method is presented to represent the stratigraphic interval as a continuous distribution by integrating over parameter uncertainty through the use of the posterior predictive distribution, which represents the stratigraphic interval as a continuous variable in time which already incorporates parameter uncertainty, rather than a bounded interval. It is possible then to combine these distributions using conflation, to build a time probabilistic model for the co-occurrence of these intervals. Empirical examples using the method for estimating the origination and extinction times of palynomorphs and the co-occurrence time for an assemblage of unknown age in the economically important Cerrejón Formation, as well as the origination time of the marine barracudas (family Sphyraenidae), are used for illustrating the potential of the method. It is implemented in the StratIntervals.jl Julia package, which allows a large set of possible prior distributions and MCMC samplers.

## 1 Introduction

There is a powerful, yet blurry window into the past of life on earth, which is the fossil record. Many insights into how the life came to be in its present form, and caveats on the possible directions that the planet can go regarding climate change and the biodiversity crisis are a few of the many applications of the knowledge that can be gathered from studying fossils (Smith et al., 2025). It has long been recognised that the fossil record is very incomplete, and multiple sources of bias make it difficult to use it for inferences about the ecosystems in deep time (Signor and Lipps, 1982). However, statistical tools have proved to be effective in dealing with the imperfections of this record, and gather insights from it through quantitative palaeobiology and its integration with evolutionary biology (Holland et al., 2024). Two of the most difficult questions to ask about any extinct lineage are when they appeared, and then when they disappeared, that is, to estimate origination and extinction times. Stratigraphic intervals have been proposed to reach this goal using information from the set time occurrences of a given lineage preserved in the fossil record, and statistical methods which describe the relationship between the pattern of such occurrences and the endpoints (i.e., origination and extinction times) of the interval (Strauss and Sadler, 1989). Although these methods were first developed with palaeobiological motivations, they have also been applied and extended in the realm of conservation biology by using collections of sightings in time as occurrences (Rivadeneira et al., 2009). In general, any system where occurrences in time are recorded, and which has start and end times could be studied using stratigraphic intervals (e.g., archaeology and cultural evolution). Furthermore, they are still useful and applicable when only one of the times is of interest, for instance, the origination time but not the extinction time, or vice versa.

There are several methods available for inference on stratigraphic intervals, with a particular focus on confidence intervals on endpoint parameters. These methods vary in assumptions (e.g., whether preservation potential is uniform), statistical paradigm (e.g., whether frequentist or Bayesian), and motivation for development (e.g., applied to palaeobiology or conservation biology). Comprehensive reviews on this subject are available in the literature (Marshall, 1990; Solow, 2005; Rivadeneira et al., 2009; Marshall, 2010; Boakes et al., 2015; McCrea et al., 2024). Rivadeneira et al. (2009) arranged the methods known at that time into three classes: Class 1 methods, which assume constant preservation potential (e.g., Strauss and Sadler, 1989; Solow, 1993a; Ferraz et al., 2003; McInerny et al., 2006), Class 2 methods, which relax that assumption (e.g., Solow, 1993b; Marshall, 1997; McCarthy, 1998; Ferraz et al., 2003), and Class 3 methods, which are distribution-free (e.g., Marshall, 1994; Solow and Roberts, 2003). Later methods, especially that of Wang et al. (2016) and the one presented here fall in Class 2 because preservation is not assumed to be constant but still rely on distributions.

Bayesian approaches have been proposed since the first attempts at estimation on stratigraphic intervals. Strauss and Sadler (1989) already recognised the advantages of Bayesian formulations of their method and provided a specific one using a uniform prior and a likelihood function constrained to the interval [0, 1]. Weiss and Marshall (1999) assume a constant preservation potential and were the first to consider discrete sampling and therefore a non-continuous likelihood function. Later, Weiss et al. (2004) extended this framework for dealing with non-constant preservation by using abundance data. Both methods are different from any other Bayesian one in that the likelihood function requires discrete sampling. Ferraz et al. (2003) provided two other Bayesian formulations, one assuming linearly decreasing preservation potential, and one uniform, in which extinction occurs suddenly. They favoured the former model, claiming that it is more realistic. Wang et al. (2016) proposed a flexible method in which different types of preservation potentials can be fitted, although they used fixed priors for the free endpoint and the preservation potential parameter. Then they used analytical formulae to calculate exactly the posterior distributions of the model parameters. However, the method conditions on one of the endpoints being zero. Alroy and Solow discussed in detail several aspects of Bayesian methods starting with a pair of papers by Alroy (2014, 2015) applied to the estimation of extinction, including the specification of priors and justification for likelihood functions (Alroy, 2016a,b; Solow, 2016a,b). Finally, Kodikara et al. (2020) proposed a method for estimating the year of extinction using sighting data and implemented the hierarchical Bayesian model to sample the posterior parameter distributions. Wang (2005) examined the correlation between parameters in Beta models and concluded that endpoint parameters tend to be correlated, which is an issue for inference using Metropolis-Hasting samplers. It is clear from the diversity of approaches using Bayesian techniques that both flexible models and their implementation in computational tools are necessary to explore and extend these Bayesian methods. The goal of the present paper is to provide a general and flexible implementation of stratigraphic intervals which allows the estimation of both origination and extinction. Also, a formulation is presented where the probability density function describing the lineage in time is not bound to concrete endpoint parameters but instead continuous, and finally a method for estimating the co-occurrence of several such lineagues in time.

## 2 The model

### 2.1 The imperfection of the fossil record and stratigraphic intervals

Consider the nature of the fossil record for a single species through time, for instance, the Ammonite species in Figure 1. It starts existing at a given point in time through speciation, stay alive for an interval of time (coloured polygon), and then for different reasons it becomes extinct. Such time interval is called a stratigraphic interval, encompassing the origination and extinction times of a species in geological time. If complete access to the history of the species was available, for instance using a time machine, it would be possible to know exactly its origination and extinction times, and its abundance through time, amongst other relevant details such as its distribution. However, the time at which species existed in the past is often well far from the present time. Also, please note that so far a time machine is yet to be developed, and is therefore unavailable. Because of the imperfect record of its existence through time, the only source of information available is in the form of fossil occurrences, which is a collection of one or more fossils at given points in time for a given species. These are samples taken from the species’ stratigraphic interval. For numerous reasons this fossil record is less than ideal for accessing basic information on the time of existence of a species in geological time.

**Figure 1:**
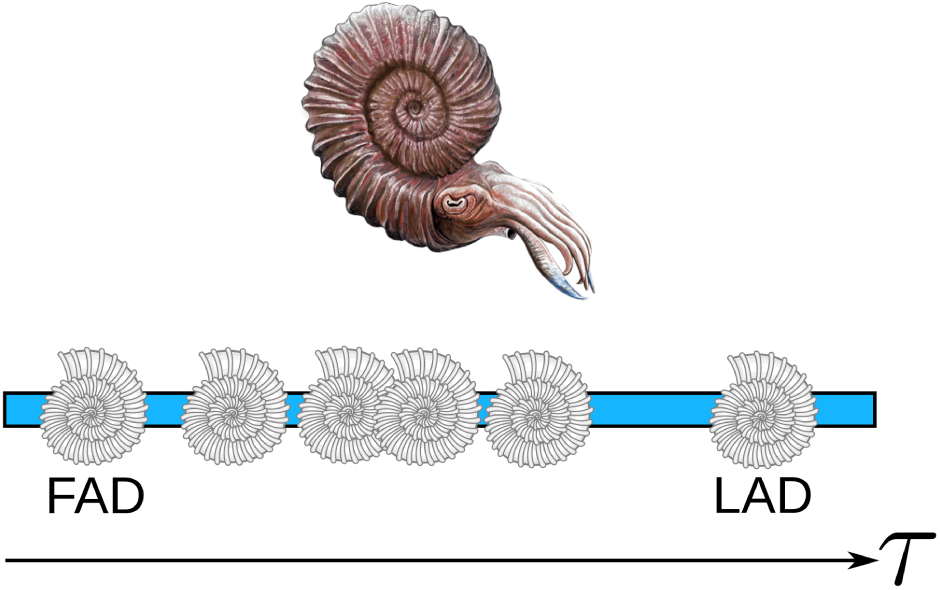
An Ammonite spanning the time interval depicted by the blue bar. Preserved fossil occurrences represented by the grey shells on the bar. The underlying distribution (the generating process) is unknown, and the only access to it are the occurrences in time themselves. First and last appearance data (FAD and LAD) are the extreme observed occurrences in time, and are always systematically younger and older than the true origination and extinction times respectively (that is, biased). Horizontal axis represents time *τ*. Ammonite reconstructions available under Creative Commons license thanks to Kouhei Futaka and Izolende.

One of the most important consequences of the imperfection of the fossil record is the so-called Signor-Lipps effect (Signor and Lipps, 1982), which states that the last occurrence of a given species in the fossil record does not match the extinction time of the species. The opposite is also true, that is, the first occurrence does not match the origination time (Figure 1). This makes sense because the probability that the first and last occurrences found exactly match the time at which the species started existing and became extinct is very very small (but never zero). These occurrences take special names in biostratigraphy: FAD and LAD, first appearance datum, and last appearance datum, respectively (Figure 1). Because of the Signor-Lipps effect and its counterpart, if the FAD and LAD were used as estimators of the origination and extinction times, it results obvious that they are biased estimators of the true times. In statistics, these are called biased because they systematically depart from the true value: The FAD is systematically younger than the origination time, and the LAD is systematically older than the extinction time. As a consequence, if FADs and LADs are used to estimate the stratigraphic interval of the species, estimates will be biased, always being shorter in magnitude than it should be in reality. Taking into account how commonplace FADs and LADs are in biostratigraphy should be concerning as these times are used for defining biozones, that is, time intervals for dating. Biostratigraphic schemes based on these are expected to be biased, being always narrower than they should be.

### 2.2 Modelling with statistics

Three important concepts were introduced in the subsection above. First, that there is some intherent probability of matching the true orgination and extinction times of a stratigraphic interval that almost never is 1.0 but on the contrary is expected to be very small. Second, that fossil occurrences in time for a given species are samples taken from the stratigraphic interval defined as the pair origination time and extinction time. Third, that if FAD and LAD are used as estimators of the true origination and extinction times, these are biased, and result in a magnitude of the stratigraphic interval which is also biased, because it is shorter than the real one. These three concepts belong to statistics, and therefore it is reasonable to use statistical methods for developing a method for estimating the origination and extinction times of a stratigraphic interval. There are three additional reasons for this. First, the availability of techniques for correcting bias in estimation. Second, statistics allows to use samples for estimating unknown quantities, called parameters. Third, the use of computing for implementing these mathematical models which help measuring uncertainty in estimation. Statistically speaking, the true but unobserved origination and extinction times in a stratigraphic interval are parameters which are to be estimated. The whole interval will now be depicted with a statistical distribution describing the relative probability of recovering fossil occurrences through time (Figure 2A). This is the statistical model, which relates the data (occurrences) as a random variable, and unknown quantities of interest, the parameters, which describe the properties of the distribution of data. In a statistical model, the data are realisations of an underlying statistical distribution with fixed parameter values (Figure 2A), and hopefully can be used to infer back the specific parameter values for such distribution. There are two important approaches in statistics: Maximum likelihood estimation, and Bayesian inference, which are introduce briefly next for a simple model with a single parameter *θ* and observations *X*. For convenience, assume that random variables and parameters do follow continuous probability distributions.

**Figure 2:**
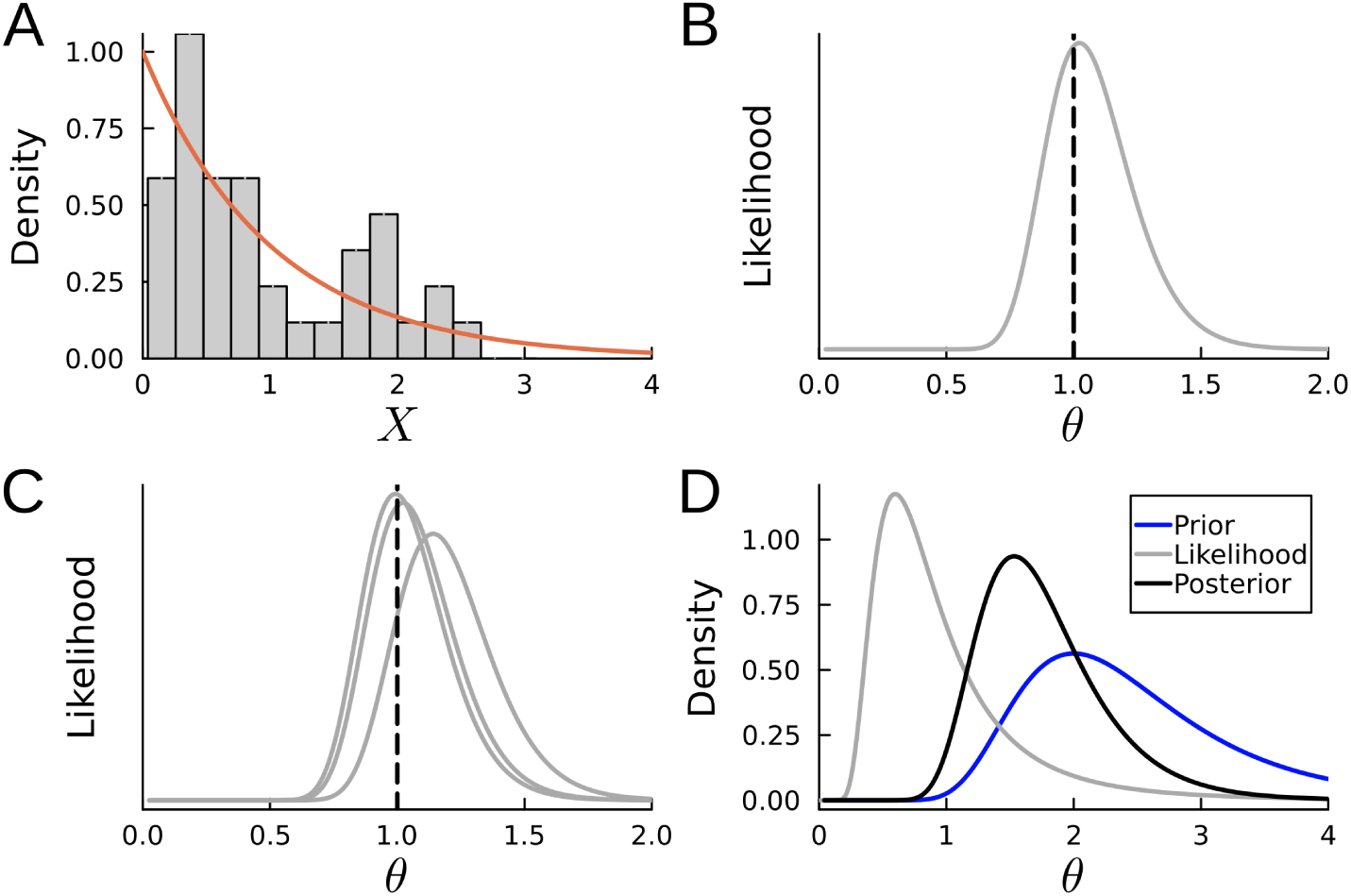
An overview of statistical foundations of the method herein developed. (A) There is an underlying statistical distribution (in orange), and the data (*X*) are realisations of if (in grey). There is an equation which relates the data *X* and the parameter(s) *θ*, which we call the statistical model, the likelihood function. (B) When fixing the data and plotting the likelihood function agains the possible parameter values, one arrives at the likelihood curve (or surface, if more than one parameter in the model, in grey). The maximum likelihood estimate is the value which maximises such curve. (C) Different datasets of the same size can generate different likelihood curves (in grey) and therefore have different maximum likelihood estimates, all of them different from the true value (black dashed line). (D) In the Bayesian framework, the likelihood function is again the statistical model, but now the use of a prior on the parameter *θ* is also used for obtaining the posterior distribution of the parameter. Note how the posterior conveys the relative information of both prior and likelihood. Note how (A) is a function of the data *X* whereas (B–D) are function of the parameter *θ*.

In maximum likelihood estimation, the statistical model *f* (*X*|*θ*), called the likelihood function, is used in numerical optimisation methods for finding the best parameter value in *θ* that maximise the fit between the observed distribution of the data *X* and the theoretical curve of the function. When the likelihood function is plotted as a function of parameter values, it is possible to perceive that maximum likelihood estimation has its name because the best parameter values are those at the maximum of the surface depicted by the likelihood function once data values are fixed (Figure 2B), as are observations which cannot change anymore. A drawback of this approach is that the likelihood function is expected to change as different data are observed, and therefore the shape which may inform about uncertainty cannot be read from the likelihood function: different data collections show different surface shapes (Figure 2C). If the surface, which is supposed to be maximised, changes with the observed data, what is the point in doing maximum likelihood estimation? Because theory says that if the data were generated by the true model, and as sample size grows to infinity, the variance of point estimates of that size will tend to zero (i.e., consistency; Lehmann and Casella, 1998). Here more data hold promise of providing better estimates.

The Bayesian framework takes on a different point of view. Here the goal is to estimate probability distributions for each of the parameters *θ* in the model instead of their point estimates, which gives access to both point estimates and uncertainty in a natural and principled way. Bayesian inference works by calculating the distribution of parameters conditional to the fixed data *X*, which is also called the Bayes’ theorem:

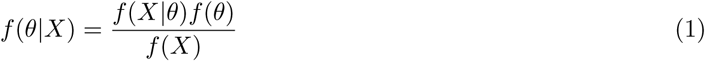

The expression above simply says that it is possible to factorise the conditional expression *f* (*θ*|*X*) so that it can be calculated using another conditional expression *f* (*X*|*θ*) multiplied by *f* (*θ*). Note that the component *f* (*X*|*θ*) is the likelihood function seen before. The expression *f* (*θ*) is the distribution of the parameter *θ* unconditioned, which is also called the prior, and is often described as the prior beliefs about possible parameter values for *θ*. More technically, the prior is just a distribution of the parameter which represents not just its point estimate but also the uncertainty around it, unconditional to the observed data *X*. Finally, the term *f* (*X*) which is the probability of the data. It is often awkward to think about the distribution of fixed data, so it is useful to represent it as a marginal distribution of the likelihood and prior with respect to the parameter *θ*. This means, that integrating over the expression *f* (*X*|*θ*)*f* (*θ*) with respect to *θ*, the parameter disappears as it is accounted for, leaving the expression as a function of *X* alone:

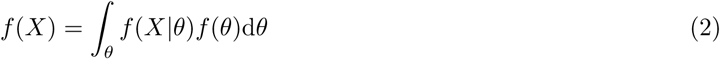

The expression above can now be used into the Bayes’ theorem. This denominator is sometimes called the normalising constant or marginal likelihood, and has the effect of guaranteeing that the expression *f* (*θ*|*X*) integrates to 1.0, which is a property of probability distributions. To conclude, *f* (*θ*|*X*) is called the posterior distribution of the parameter *θ*:

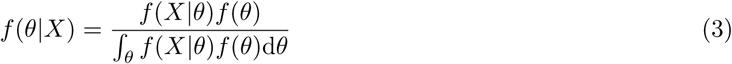

An important property of the expression above is that the posterior distribution of *θ* is defined by both the likelihood function and the prior proportional to their variances. Also, variance is inversely proportional to information (and directly proportional to the Shannon entropy, Shannon, 1948; Ebrahimi et al., 1999), and therefore the posterior will tend to be closer in values to whatever distribution has the lowest variance, or well in the middle if both distributions have the same variance. This effect can be seen in Figure 2D for fixed the data *X* and then plotting the conditional distribution with varying *θ* for the prior, likelihood, and posterior distributions. Note that this plot is not necessarily equal to the curve *f* (*X*|*θ*), which is a primary function of *X*. In conclusion, the Bayesian approach ponders the relative information content in both the likelihood (data) and the prior. Quite often, it any previous guess of the possible distribution of parameter values is simply unavailable, it is still possible to use what is called wide, flat, or weakly informative priors, which have high variance and therefore should allow the posterior to be more informed by the likelihood rather than the prior.

If the expression *f* (*θ*|*X*) is algebraically simple, and if 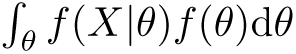 can be calculated analytically, then a closed formula can be found for the posterior distribution of *θ*. With this, probabilities of observing specific intervals of parameter values (e.g., *P* (*X <* 0.1), *P* (0 ≤ *X* ≤ 10)) can be calculated. Otherwise, it is necessary to use Markov Chain Monte Carlo (MCMC) algorithms for sampling from these posterior distributions without need of closed formulae, by approximating them with finite samples. MCMC algorithms are popular because analytical versions of posterior distributions are very rare in but the simplest models, or in those that use conjugate priors (Lindley, 1972).

Once the posterior distributions of the parameters are available, a new distribution of the random variable which is a function of the observed data and is marginalised over the parameter uncertainty can be constructed. This is called the posterior predictive distribution and can be described as a distribution of the data taking into account the uncertainty in the parameters from the posterior distribution. The term posterior predictive comes from the fact that such marginalised distribution can be used either for calculating the probability of observing concrete future data, or for simulating future data. These distributions are useful in many modelling applications but rarely used in evolutionary biology. This is accomplished with the following integration:

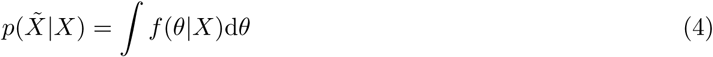

The posterior predictive it thus a distribution that no longer depends on the parameters as these have been integrated out and incorporated in the uncertainty around (future) values of 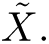 This integration can be carried out with a posterior sample of parameter values conditional to the observed data *X* and therefore its implementation after use of MCMC algorithms is straightforward. The use of a finite posterior sample produces also a finite posterior predictive sample of 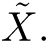 This can be approximated as a continuous function by an interpolated univariate kernel density (Silverman, 1998), using a vector of posterior predictive samples as input. This produces a continuous function of 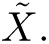

### 2.3 Likelihood function of a stratigraphic interval

A stratigraphic interval (Strauss and Sadler, 1989) is a segment of time bound by the origination and extinction times, where a lineage is existing on earth. Each stratigraphic interval represents the known occurrences of a given lineage in geologic time. These lineages have unobserved origination and extinction times, parameters *θ*_1_ and *θ*_2_ respectively (Figure 3A). The occurrences of the lineage are preserved in the fossil record in a way which has been described by different models in the literature, from uniform to distribution-free ways. It is reasonable to consider that preservation is most likely non-uniform across geologic time for a given lineage, but that it may also be uniform. This can be accomplished by using a generalised beta distribution which has a very flexible set of shapes governed by the parameters *α* and *β*, which jointly describe the shape of the distribution of occurrences in time. Together these can describe both non-uniform and uniform preservation patterns. A reparametrisation in order to describe an comparably rich set of preservation possibilities is applied here in order to reduce the free parameters in the model while retaining flexibility.

**Figure 3:**
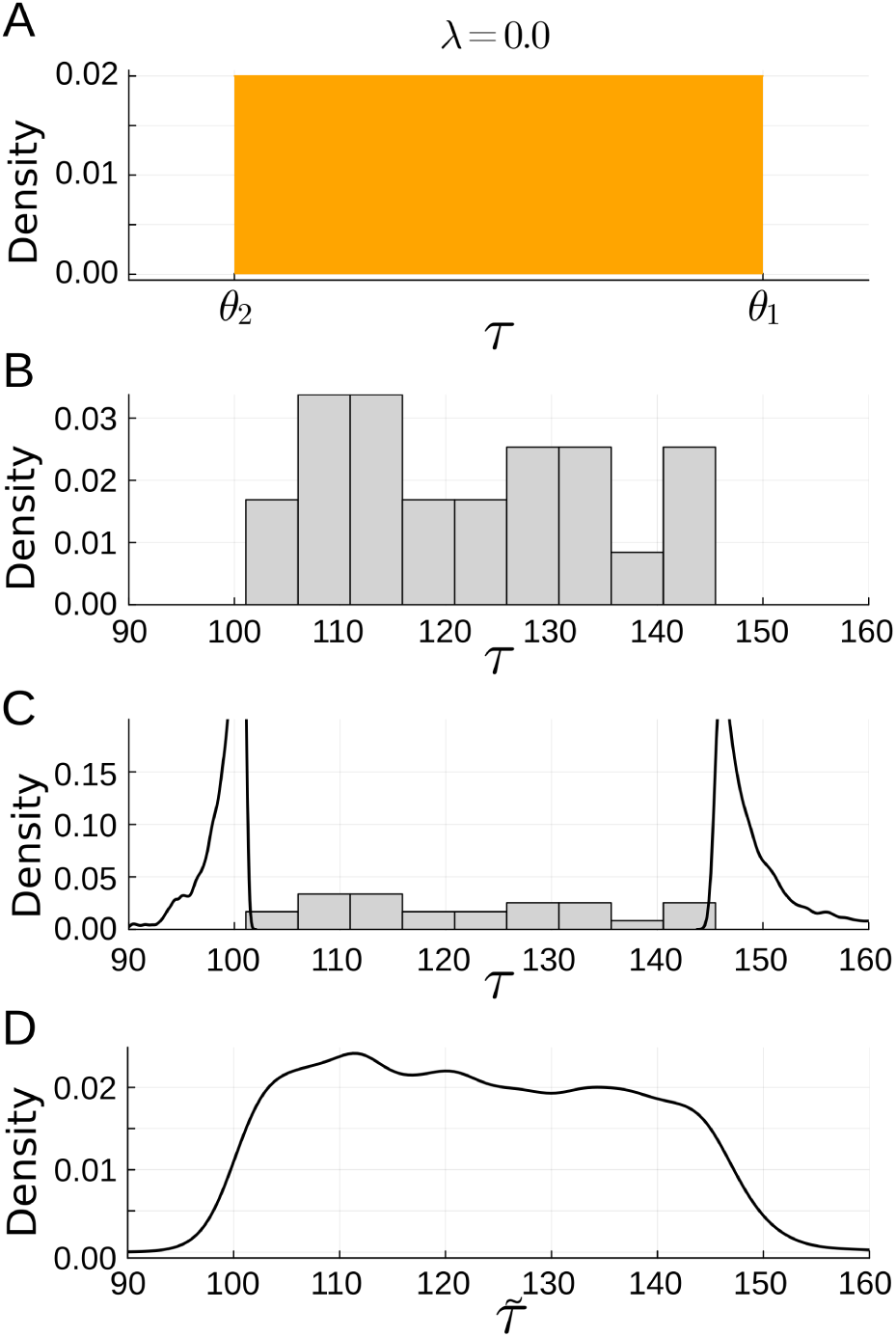
A stratigraphic interval (A) is defined by the underlying distribution *f* (*τ* |*θ*_1_*, θ*_2_*, λ*) (in orange, uniform in this example), whose probability density function is defined by the recovery potential governed by *λ* with time occurrences [*τ*_1_*,…,τ*_6_], which are realisations of said distribution, as seen in (B). The posterior distributions of *θ*_1_ and *θ*_2_ are then approximated using MCMC sampling algorithms, here plotted as the density plots depicting the regions of time where such parameters are expected to be (C). Note that such distributions provide measures such as means and medians for summarising the centre of the distributions, as well as uncertainty, for instance using the highest posterior density interval. When conditioning to the observed occurrences and integrating over parameter uncertainty, one arrives at the posterior predictive distribution, which is a continuous function of *τ* that already accounts for parameter uncertainty. This is the continuous form for representing the life of a lineage in geological time, and no longer needs to be bound to concrete values of *θ*_1_ and *θ*_2_.

The data in this model are the fossil occurrences in time. Let *τ* be a random variable representing an occurrence of this type. Although in principle such interval in continuous, only point time occurrences inside the bound interval are available or can be sampled, whose distribution is governed by a parameter or set of parameters describing the potential of preservation (Figure 3B). A function that describes the likelihood of observing a given time occurrence with the four-parameter Beta distribution (e.g., Wang, 2005) defined over *τ* ∈ [*θ*_1_*, θ*_2_] is constructed as:

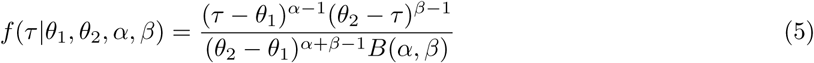

where *B*(*α, β*) is the Beta function.

Now reparametrise the function so that it depends only on a parameter *λ* instead of *α* and *β* in a similar way as in Wang et al. (2016). This has the advantage of requiring less data while still giving the likelihood function enough flexibility to describe different preservation scenarios (Figure 4). This function is called the Three-Parameter Beta distribution. Note however that here none of the endpoint parameters are fixed, which gives the likelihood function:

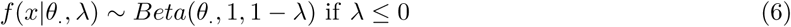

**Figure 4:**
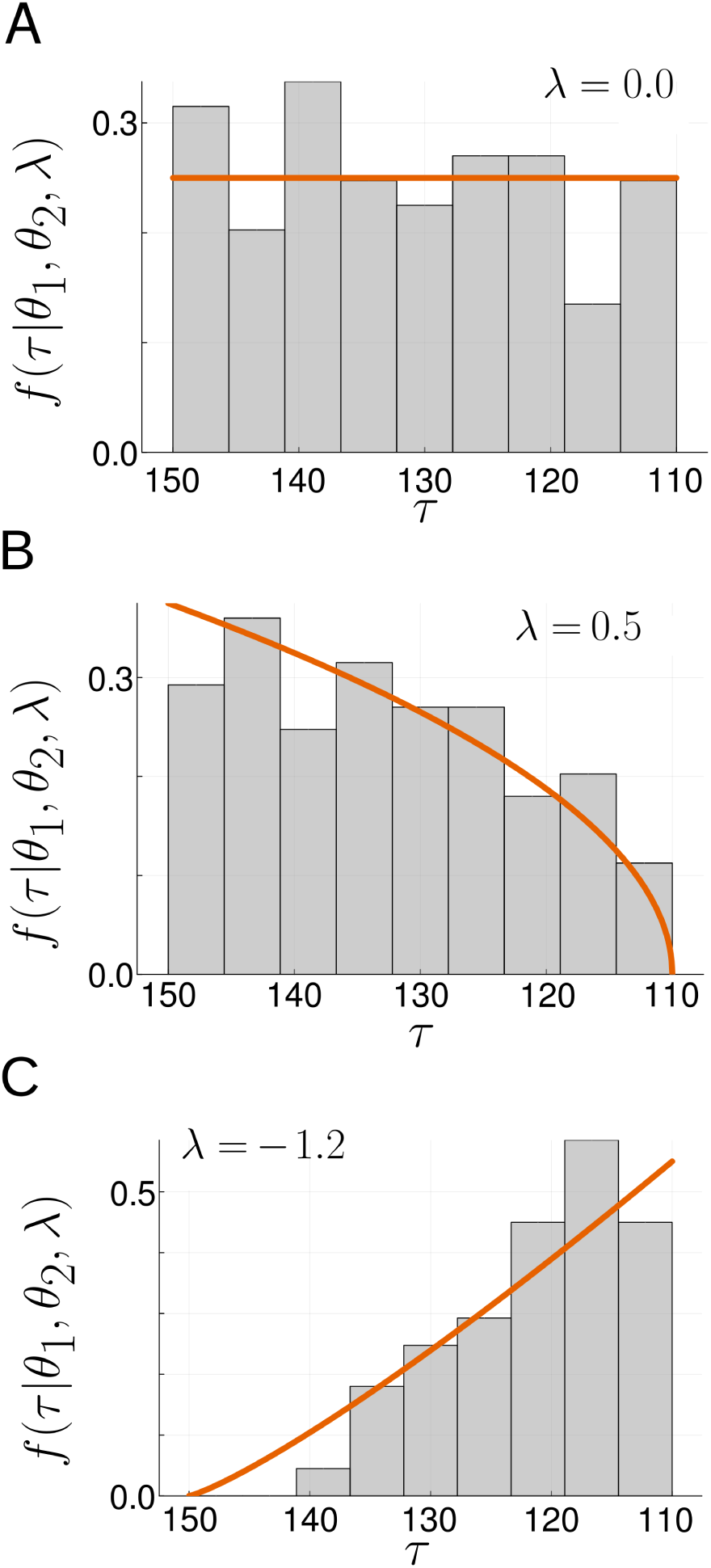
(A) When *λ* = 0.0, the distribution of occurrences is uniform through time. (B) When it is negative the trend is decreasing in time, and (C) the opposite when it is positive. The independent variable is time, depicted by *τ* in all figures, whereas the dependent variable is the probability density function *f* (*τ* |*θ*_1_*, θ*_2_*, λ*). The curve in orange depicts the true underlying distribution, and the grey histogram represents a representative realisation.

and

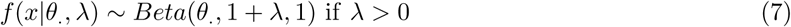

which gives

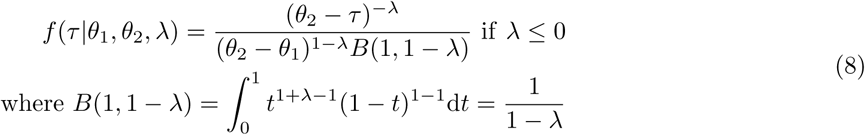

and

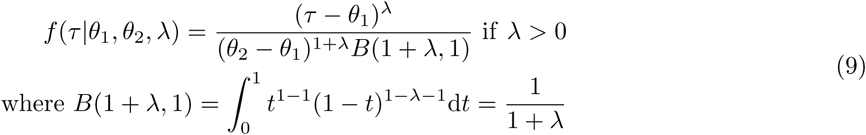

Assuming that time occurrences are independent and identically distributed, the likelihood of observing a vector representing the collection of *N* time occurrences given a stratigraphic interval is calculated with the product of likelihoods of individual time occurrences:

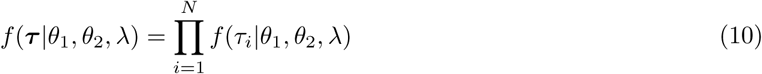

The model above has quite some flexibility in the types of shapes that takes depending on the value of *λ*. When *λ* = 0.0 the distribution becomes uniform, in the same way as a the Four-Parameter and Standard Beta do when both *α, β* = 1.0 (Wang, 2005). The distribution takes a triangular shape when *λ* = 1.0, and it does have an asymmetric shape which is convex towards the tail on *θ*_2_ when 0.0 *< λ <* 1.0 and, and concave when *λ >* 1.0. The position of the tail just shifts to *θ*_1_ when *λ* is negative in these same intervals.

The behaviour of flat likelihood curves which still have a maximum at their observed minimum and maximum data values when their likelihood surfaces are plotted as a function of parameter values instead of the data is also seen in other implementation of uniform likelihood functions (e.g., see Delignette-Muller and Dutang, 2015). It is therefore concluded that such parameters are identifiable, but biased. The reason is that the likelihood for the endpoint parameters is problematic in non-regular models, which are those where the support of the likelihood function depends on paramaters (Sareen, 2003), as is the case of the Three-Parameter Beta. Several authors have pointed out to the issues when trying to use maximum likelihood estimation in non-regular models (Wang, 2005; Hall and Wang, 2005; Sareen, 2003). All these authors however point to the use of priors to constrain the estimation of endpoint parameters as the best alternative available for the estimation of these model parameters. A Bayesian model is herein defined using the Three-Parameter Beta as the likelihood function.

### 2.4 The posterior distribution of parameters and posterior predictive of *τ*

The Bayes’ theorem is used in order to construct the posterior densities for these parameters given the observed vector of occurrences in time (Figure 3C):

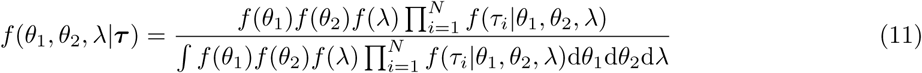

Depending on the specific choice of priors for *θ*_1_, *θ*_2_, and *λ*, this expression may have a closed form (e.g., using conjugate priors). However, for several prior functions this expression may not be closed and therefore MCMC algorithms can be used for sampling from the posterior distribution even when the integral in the denominator cannot be calculated analytically.

Now that the posterior parameter distributions are available, it is possible to sample from this conditional distribution in order to build a posterior predictive distribution of *τ* by integrating over the parameter uncertainty. This posterior predictive distribution represents the probabilistic model for the occurrences through time while accounting for uncertainty in endpoint and preservation parameter estimation (Figure 3D).

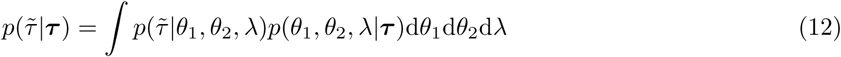

### 2.5 The distribution of the co-occurrence of multiple stratigraphic intervals

Suppose that multiple lineages are present in a given fossil assemblage, that is, a fossil community composed of different species which coexist in space and time. Each of these species is expected to be also known from other points in time (and maybe space) and thus each of the species can be represented by a set of occurrences *τ* composed of all the points in time at which it is known that the lineage existed (e.g., the shells representing fossilised time occurrences in Figure 5). The goal here is to estimate the time of co-occurrence, that is, a distribution that describes when all these lineages were living together, by combining the distributions of each stratigraphic interval.

**Figure 5:**
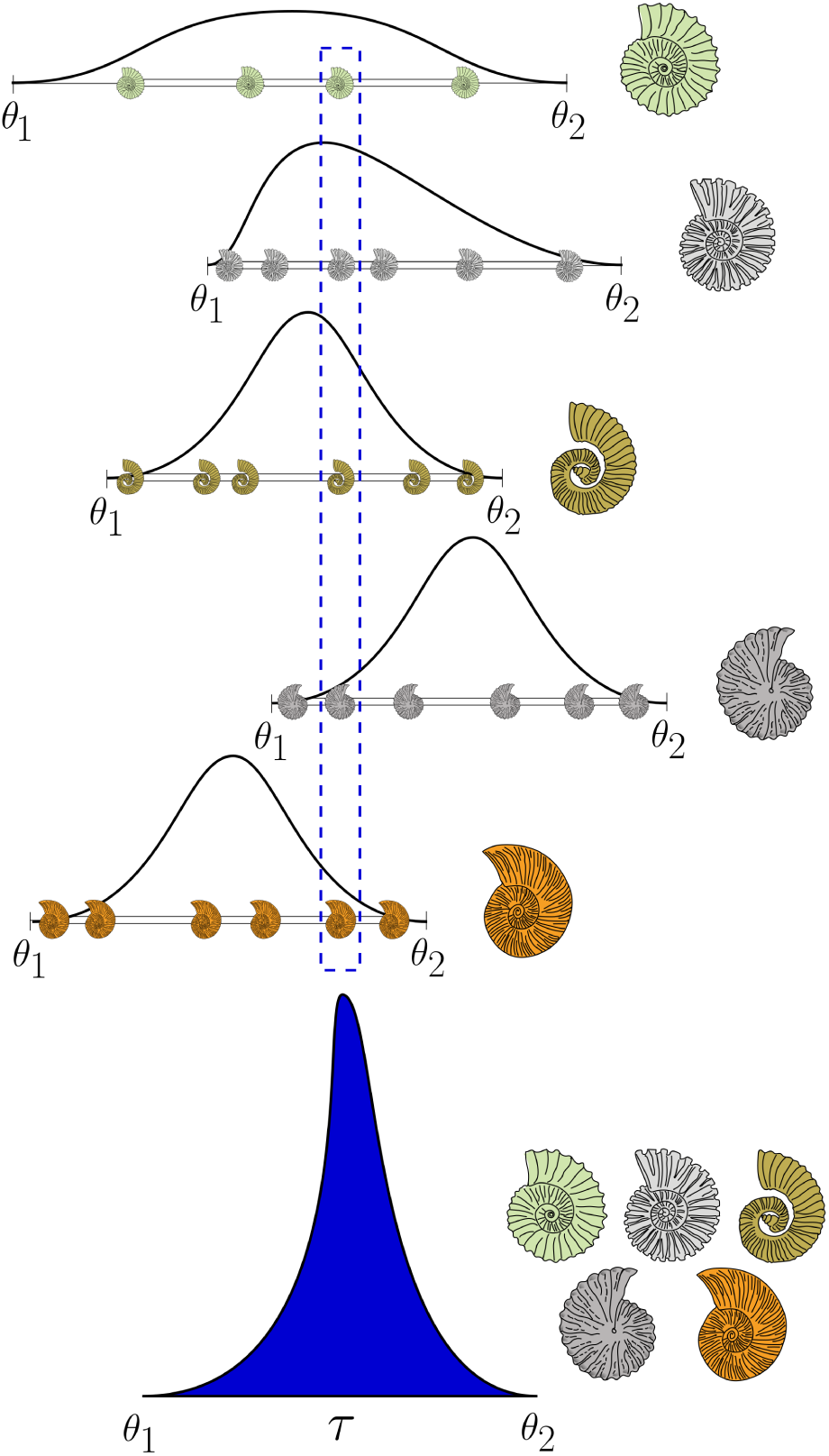
Assemblage generating process. Each lineage has an origination and extinction time designed by *θ*_1_*, θ*_2_. Coloured ammonites represent occurrences in time, and assuming that all except those inside the blue dashed region have their age given, so that the goal is to estimate their age in that region, which represent the coexistence time of all five lineages. Such time is given by the conflation (blue distribution below) of all the probability density functions (black lines in top four panels) representing the stratigraphic interval of each species. Ammonite reconstructions available thanks to Jorge W. Moreno-Bernal.

Combining different distributions is not straightforward (Genest and Zidek, 1986). However, the conflation of probability density functions is a useful procedure which combines them provided that each of them is independent (Hill, 2011; Hill and Miller, 2011). Assuming that each distribution describing the posterior predictive distribution of each interval is independent, and simplifying notation so that 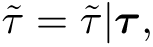 it is possible to define the composite distribution of 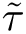 for *M* intervals as the conflation of individual posterior predictive distributions:

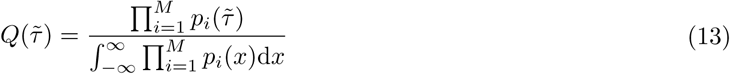

Such distribution is useful e.g. for building credible intervals for the time *τ* of co-occurrence of stratigraphic intervals. The conceptual interpretation is as follows: *M* different lineages as represented by their stratigraphic intervals should coexist at most for some time interval when they all were alive. As the conflation of densities is a density itself, it can be used for asking questions on the probability of co-existence of lineages during some arbitrary time interval given the distribution.

### 2.6 Implementation

The Bayesian formulation of stratigraphic intervals with the Three-Parameter Beta distribution as a model has been implemented in the StratIntervals.jl package, written in pure Julia (Bezanson et al., 2017). The package uses several dependencies (DataFrames, Distributions, Interpolations, KernelDensity, QuadGK, Random, SpecialFunctions, and StatsPlots; Bouchet-Valat and Kamiński, 2023; Besançon et al.,

2021; Johnson, 2013) which provide data structures, plotting engines, or implement specific mathematical and statistical procedures not available in the Base Julia library. In particular, StratIntervals uses Turing (Ge et al., 2018), a package providing a probabilistic modelling language which allows for flexible implementation of MCMC algorithms.

A generalisation of the Exponential distribution, called herein the Reflected-Offset Exponential (RefOff-Exp or RefOffExponential) with parameters *s, o, ρ* where *s* is the scale, *ρ* the reflection constant (either 1 or-1) and *o* is the offset, has been implemented in order to allow Exponential priors which start from a point different from zero, as well as a reflection parameter which allows it to decrease towards the left instead of the right in the standard Exponential distribution. The Reflected-Offset Exponential distribution has probability density function:

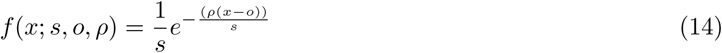

The stable version of StratIntervals can be found in the official Julia Registry. The development version can be found at https://github.com/gaballench/StratIntervals.jl. A static website with documentation and vignettes can be found at https://gaballench.github.io/StratIntervals.jl.

## 3 Simulations

### 3.1 Implementation evaluation

Three sets of simulations were performed to verify the properties of the implementation under increasing sample size and different prior specifications. For all simulations, a true interval with *θ*_1_ = 150.0 Ma and *θ*_2_ = 100.0 Ma was chosen, and a true preservation parameter *λ* = 0.0. All simulations were implemented in pure Julia code, using GNU parallel (Tange, 2011, 2024) when appropriate, and run in the server gymnotus at the Instituto de Biociências, Universidade Estadual Paulista, Botucatu.

#### 3.1.1 Maximum likelihood estimation

Increasing datasets of 10, 50, 100, 1000, and 10000 occurrences were simulated to plot the joint likelihood surface for combination of (*θ*_1_*, λ*), (*θ*_2_*, λ*), and (*θ*_1_*, θ*_2_), while keeping the remaining parameter fixed to the true value. The true value of each parameter was then compared to the likelihood surface to see if parameter estimation is consistent.

#### 3.1.2 Bayesian inference estimation

Increasing sample size of 10, 50, 100, 200, and 500 occurrences were simulated under the true stratigraphic interval. Then, MCMC sampling was used to calculate the posterior distributions of all three parameters. The no-U-turn sampler (NUTS) and the Hamiltonian Monte Carlo sampler show advantages in situations where parameters may be correlated as appears to be the case of Beta models (Wang, 2005), therefore 10000 iterations were used in the NUTS (Hoffman and Gelman, 2014) as it required to set less quantities than the Hamiltonian Monte Carlo.

The interest here is in estimating endpoint parameters *θ*_1_*, θ*_2_, and following the results suggesting an appropriate behaviour of *λ* estimation during simulations and maximum likelihood estimation, it was fixed to its true value instead.

Normal priors *N* (*µ, σ*) were set in four different combinations: Correct–incorrect, and informative–poorly-informative. A correct prior is defined as one centred at the true parameter value, whereas an incorrect prior is centred elsewhere and outside the stratigraphic interval (e.g. 20 units away from the true value). An informative prior (i.e., one with low variance) was set to *σ* = 1.0, whereas a poorly-informative one to *σ* = 20.0. Thus, for *θ*_1_ the four combinations of priors are *N* (150.0, 1.0), *N* (150.0, 20.0), *N* (150.0 + 20.0, 1.0), and *N* (150.0 + 20.0, 20.0), whereas for *θ*_2_ these are *N* (100.0, 1.0), *N* (100.0, 20.0), *N* (100.0 − 20.0, 1.0), and *N* (100.0 − 20.0, 20.0).

The effect of co-estimating *θ*_1_ and *θ*_2_ was assessed by running the simulations with priors set as described above, whereas the effect of estimating just one at a time was measured by fixing the other parameter to its true value and sampling just one at a time.

A total of 1000 simulations were run for each of these eight combinations: prior correctness, informativeness, and co-estimation.

A separate set of simulations was carried out for assessing the impact of the value of *λ* on the estimation of endpoint *θ* parameters. Here, a dataset of 75 occurrences was simulated from the Three-Parameter Beta distribution with the true value of *λ* varying (-1.5,-1.0,-0.5, 0.0, 0.5, 1.0, 1.5), whereas the true value of *θ*_1_ and *θ*_2_ were set to 550.0 and 500.0 Ma respectively, for a total of 7 simulated datasets. Priors on model parameters were set to *θ*_1_ ∼ *N* (*µ* = 550.0*, σ* = 10.0), *θ*_2_ ∼ *N* (*µ* = 500.0*, σ* = 10.0) and *λ* ∼ *N* (0.0, 2.0). The posterior distributions of model parameters were sampled using the NUTS with 100000 iterations.

### 3.2 Implementation and parameter identifiability

#### 3.2.1 Asymptotic properties and parameter estimation under maximum likelihood and Bayesian inference

The likelihood surface was found to be flat with respect to the endpoint parameters, whereas it it well-behaved for the preservation parameter. The endpoint parameter estimators are biased, which means that the first appearance datum (FAD) and last appearance datum (LAD), instead of the true value, always maximise the likelihood surface. As n rises and the LAD and FAD tend to approach the true endpoint values, and therefore the MLE converges to the true value (Figure 6).

**Figure 6:**
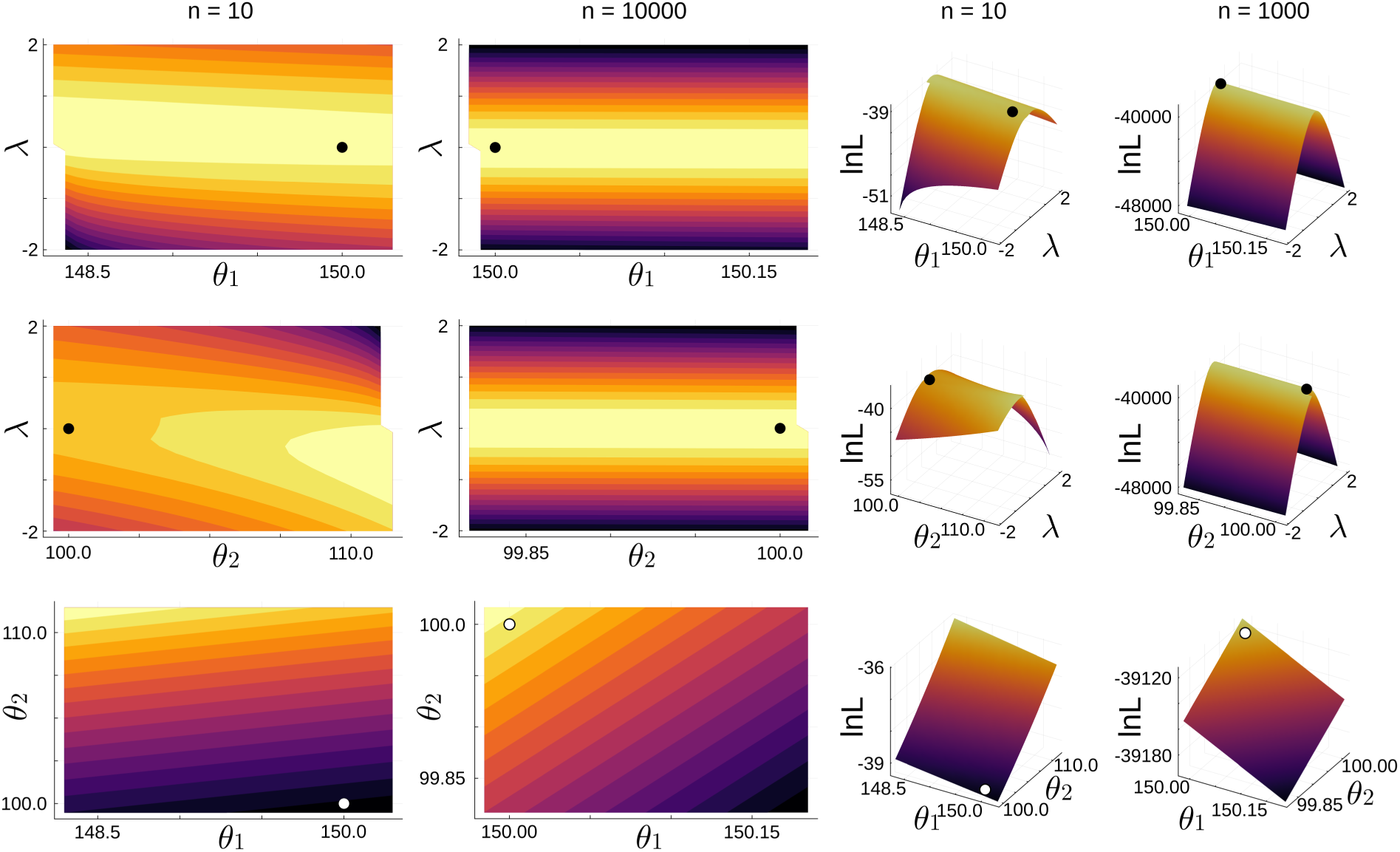
Log-likelihood surface of different pairs of parameters from the ThreeParBeta(*τ*; *θ*_1_*, θ*_2_*, λ*) distribution as a function of a fixed dataset of varying size (*n* = 10 or *n* = 10000). Left-most columns are 2-dimensional representations of the 3-dimensional surfaces to the right. Isoclines represent log-likelihood, the brighter the higher. The *x* and *y* axes are the parameter pairs (*θ*_1_*, λ*), (*θ*_2_*, λ*), and (*θ*_1_*, θ*_2_) from top to bottom. In all cases, the remaining parameter was fixed to its true value. The pair of true values is plotted as a dot, which changes in colour just to aid in visualisation. The estimation of *λ* is unbiased, whereas the estimation of both *θ*_1_ and *θ*_2_ is biased and only converges to the true value asymptotically.

The Bayesian version of the method, however, does not show bias in estimation and is better behaved in terms of estimation accuracy (Figure 7). The cost of co-estimating the both endpoint parameters in simulations as compared to fixing one of them to the true value is negligible (Supplementary Figure S1) and therefore only simulations estimating one endpoint parameter while fixing the other to the true value are herein discussed.

**Figure 7:**
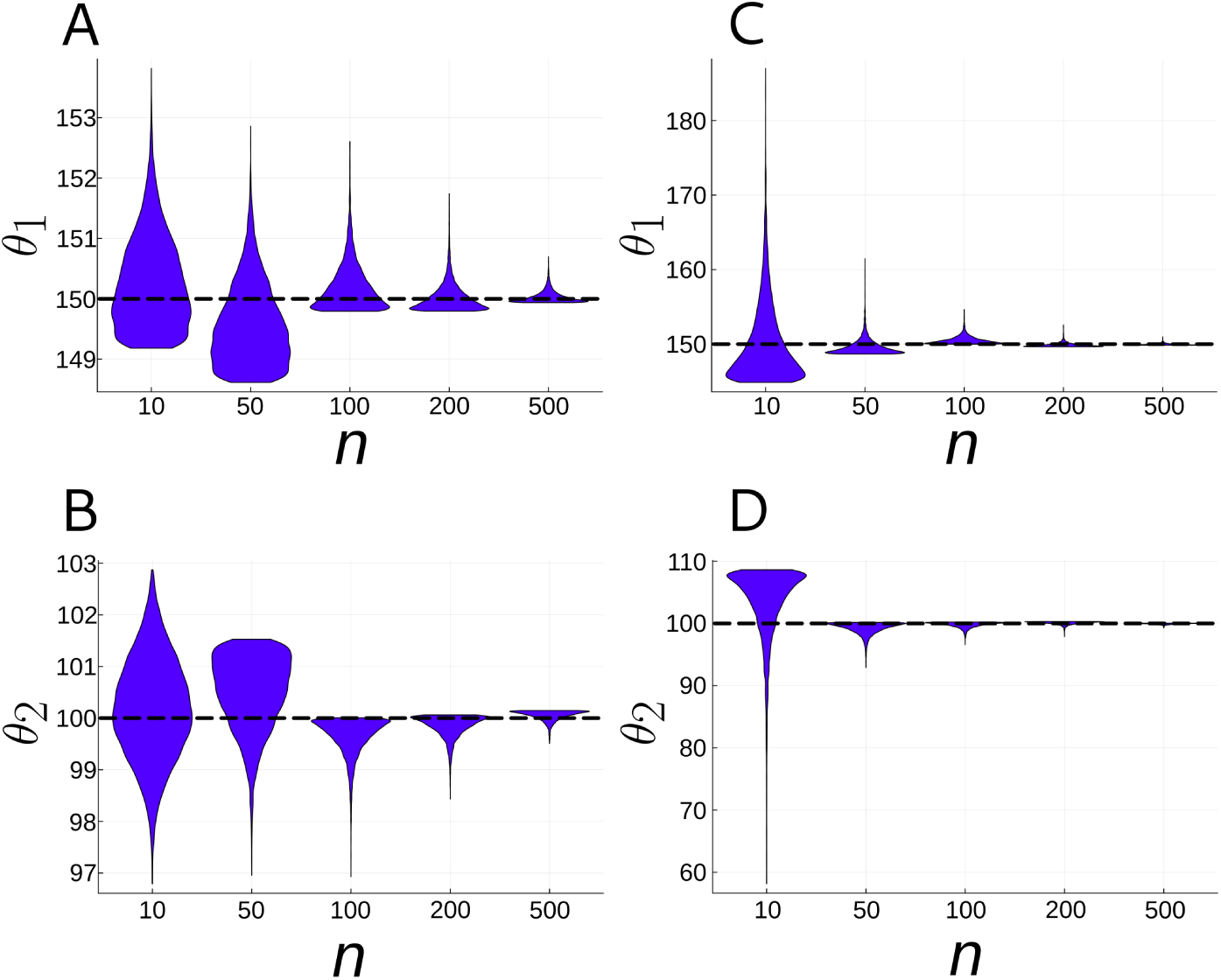
Posterior distribution of a single simulated dataset of varying sample size from the ThreeParBeta(*τ*; *θ*_1_*, θ*_2_*, λ*) distribution to illustrate the effect of sample size (*x* axis) and prior variance on *θ*_1_ and *θ*_2_ (*y* axis, in Ma). A) Posterior distribution of *θ*_1_ with prior *N* (*µ* = 150.0*, σ* = 1.0). B) Posterior distribution of *θ*_2_ with prior *N* (*µ* = 100.0*, σ* = 1.0). C) Posterior distribution of *θ*_1_ with prior *N* (*µ* = 150.0*, σ* = 20.0). D) Posterior distribution of *θ*_2_ with prior *N* (*µ* = 100.0*, σ* = 20.0). The dashed line represents the true value of the parameter in all plots.

When simulating multiple datasets (1000) of varying sample size, the distribution of posterior medians was found to concentrate around the true value (Figure 8), and the variance of this sample of medians decreases as with an increase in sample size. This measures the precision of estimation of the model in general, over multiple simulated datasets.

**Figure 8:**
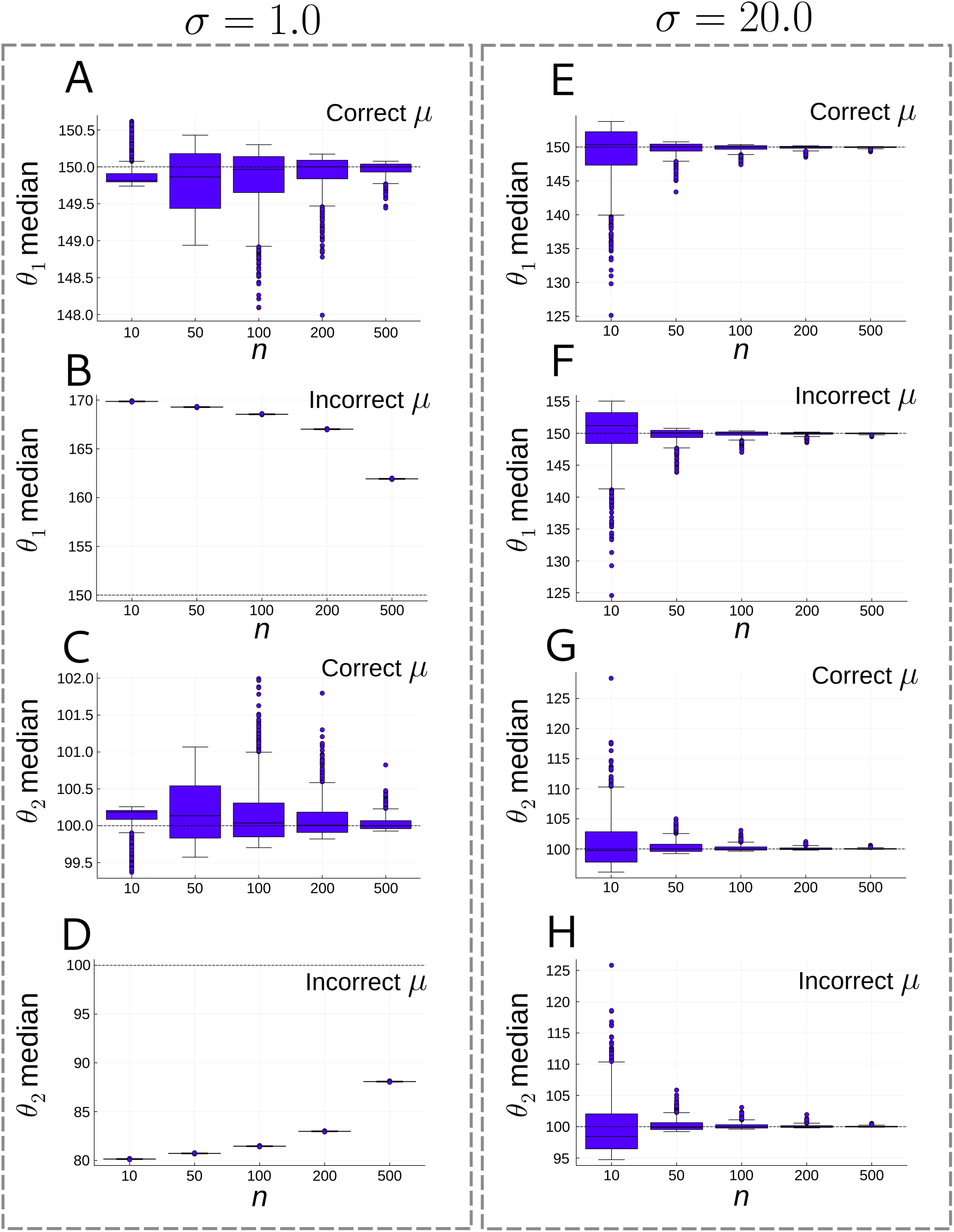
Distribution of posterior medians (*y* axis, in Ma) across 1000 simulations for combinations of correct and incorrect prior, and informative and poorly-informative prior, as a function of sample size (*x* axis). A) Correct and informative prior (*θ*_1_ ∼ *N* (150.0, 1.0)). B) Incorrect and informative prior (*θ*_1_ ∼ *N* (150.0 + 20.0, 1.0)). C) Correct and informative prior (*θ*_2_ ∼ *N* (100.0, 1.0)). D) Incorrect and informative prior (*θ*_2_ ∼ *N* (100.0−20.0, 1.0)). E) Correct and poor2ly1-informative prior (*θ*_1_ ∼ *N* (150.0, 20.0)). F) Incorrect and poorly-informative prior (*θ*_1_ ∼ *N* (150.0 + 20.0, 20.0)). G) Correct and poorly-informative prior (*θ*_2_ ∼ *N* (100.0, 20.0)). H) Incorrect and poorly-informative prior (*θ*_2_ ∼ *N* (100.0 − 20.0, 20.0)). The dashed line represents the true value in all cases.

The inter-quartile range (IQR) of the posterior distribution, which measures better the spread in asymmetrical distributions, also reduces as sample size grows (Figure 9), therefore showing a gain in precision within each posterior estimation. This measures the precision of estimation conditional to a specific dataset. However, when a prior is incorrect and informative, the precision actually decreases with sample size (Figures 9B,D). This is not the case for incorrect–poorly-informative, correct–informative, and correct–poorly-informative priors, all of which show an increase in precision with increase in sample size.

**Figure 9:**
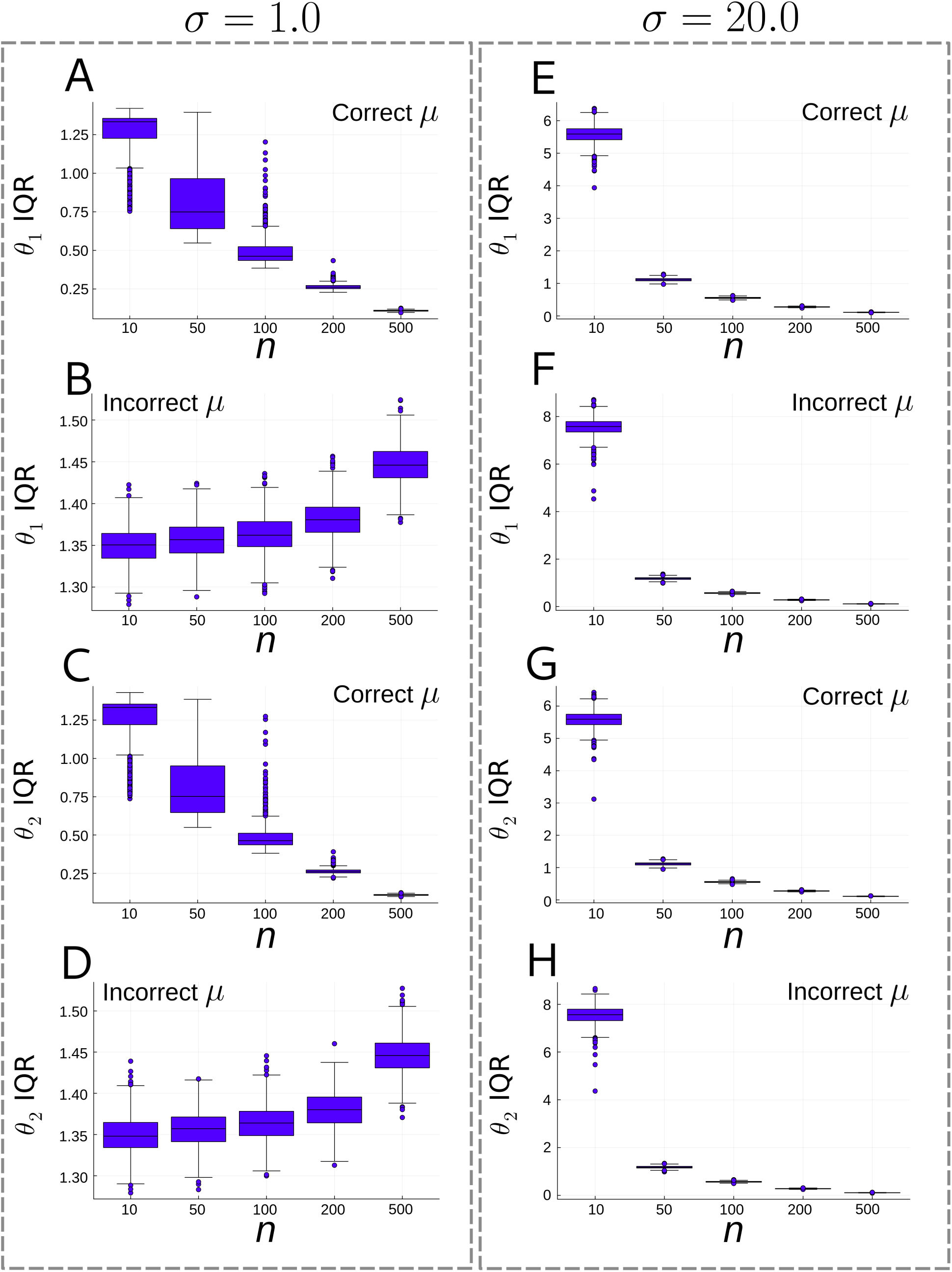
Distribution of posterior inter-quartile range (IQR, y axis) across 1000 simulations for combinations of correct and incorrect prior, and informative and poorly-informative prior, as a function of sample size (x axis). A) Correct and informative prior (θ1 ∼ N(150.0, 1.0)). B) Incorrect and informative prior (θ1 ∼ N(150.0 + 20.0, 1.0)). C) Correct and informative prior (θ2 ∼ N(100.0, 1.0)). D) Incorrect and informative prior (θ2 ∼ N(100.0−20.0, 1.0)). E) Correct and poorly-informative prior (θ1 ∼ N(150.0, 20.0)). F) Incorrect and poorly-informative prior (θ1 ∼ N(150.0 + 20.0, 20.0)). G) Correct and poorly-informative prior (θ2 ∼ N(100.0, 20.0)). H) Incorrect and poorly-informative prior (θ2 ∼ N(100.0 − 20.0, 20.0)).

The mean squared error (MSE) lowers by raising sample size (Figure 10), which is consistent with the results of the IQR, and highlights both the increase in accuracy and precision during estimation across multiple simulated datasets from the same model. The decrease in MSE is consistent regardless of the presence of incorrect and/or poorly-informative priors.

**Figure 10:**
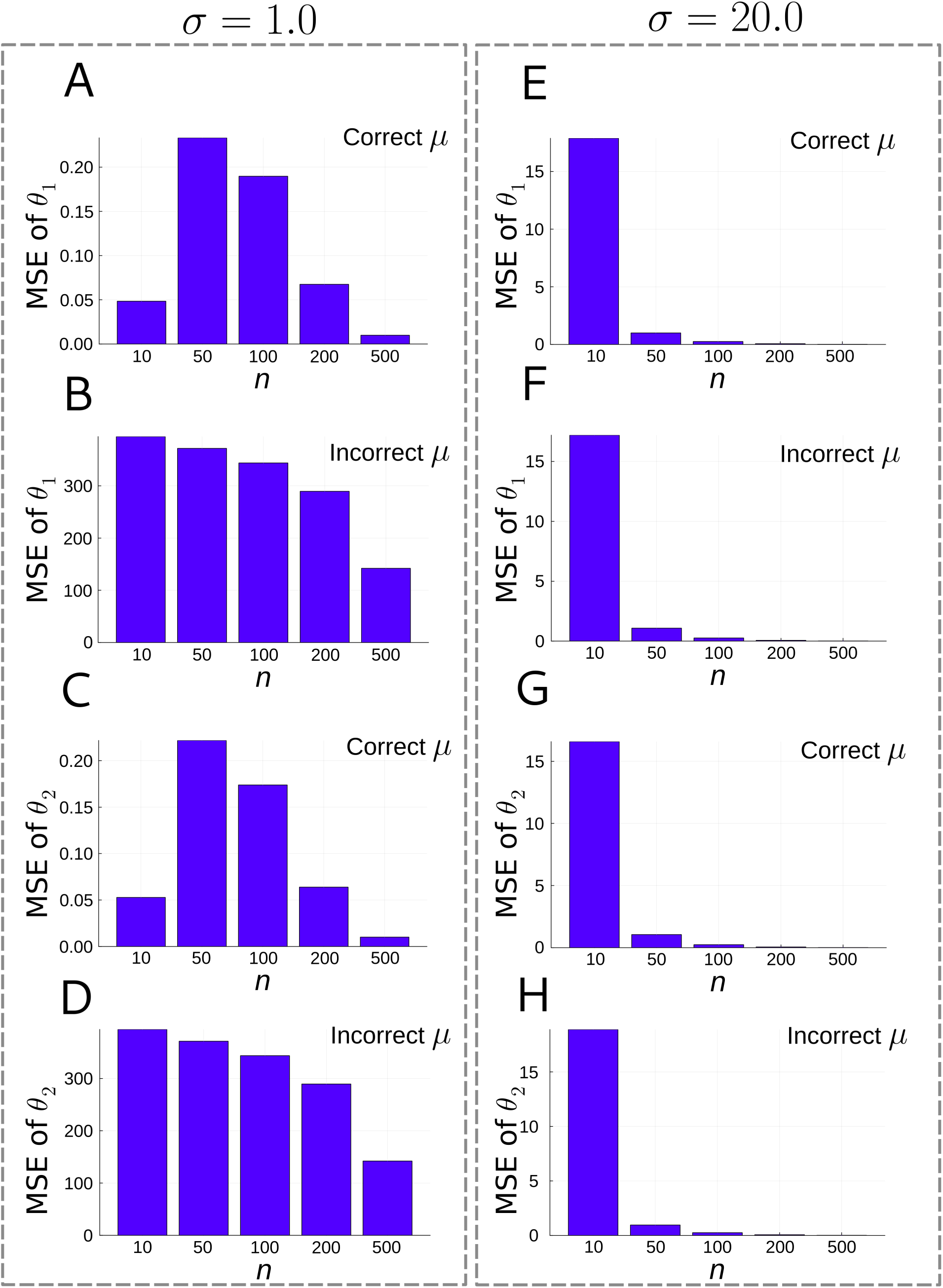
Mean squared error (MSE, y axis) across 1000 simulations for combinations of correct and incorrect prior, and informative and poorly-informative prior, as a function of sample size (x axis). A) Correct and informative prior (θ1 ∼ N(150.0, 1.0)). B) Incorrect and informative prior (θ1 ∼ N(150.0+20.0, 1.0)). C) Correct and informative prior (θ2 ∼ N(100.0, 1.0)). D) Incorrect and informative prior (θ2 ∼ N(100.0 − 20.0, 1.0)). E) Correct and poorly-informative prior (θ1 ∼ N(150.0, 20.0)). F) Incorrect and poorly-informative prior (θ1 ∼ N(150.0 + 20.0, 20.0)). G) Correct and poorly-informative prior (θ2 ∼ N(100.0, 20.0)). H) Incorrect and poorly-informative prior (θ2 ∼ N(100.0 − 20.0, 20.0)).

The value of *λ* and therefore the shape of the Three-Parameter Beta distribution have an influence on the posterior variance of the *θ* endpoint parameters (also see Supplementary Figure S2). When *λ* = 0.0, and therefore the model has a uniform shape, the posterior variance of both *θ*_1_ and *θ*_2_ is the same (Figure 11A). However, when *λ* is positive (e.g., 1.5), the tail goes to the present (Figure 11B), whereas if it is negative (e.g., =-1.5), there is a tail towards the past (Figure 11C).

**Figure 11:**
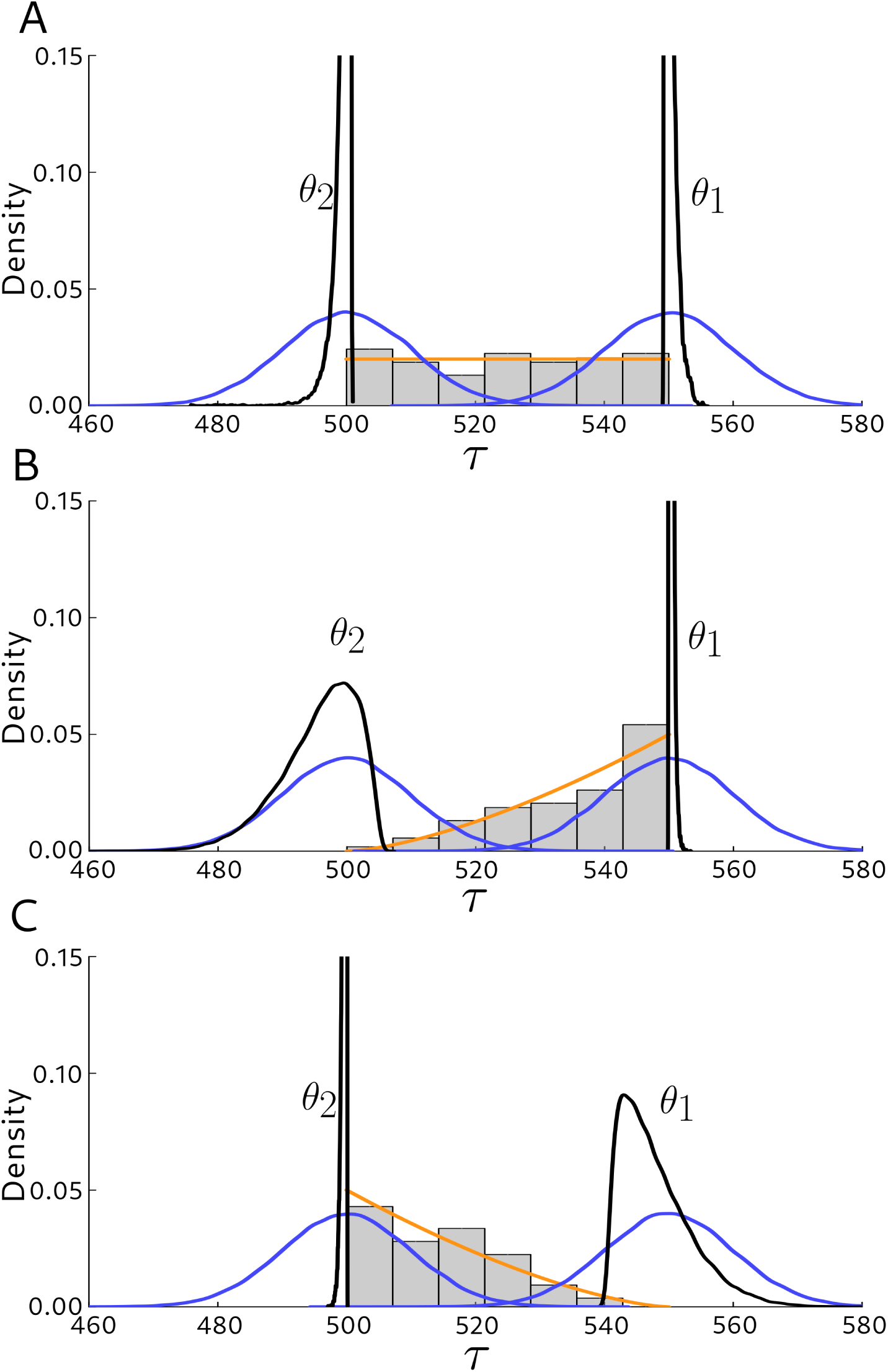
Effect of *λ* on *θ*_1_ and *θ*_2_ for a sample size of 75 occurrences sampled from distributions with different values of *λ*. A) *λ* = 0.0. B) *λ* = 1.5. C) *λ* = −1.5. In all plots the orange line represents the ThreeParBeta(*τ*; *θ*_1_ = 550.0*, θ*_2_ = 500.0*, λ*) from where the 75 occurrences were sampled. Blue lines represent the rather poorly-informative prior centred at the true value on each (*θ*_1_ ∼ *N* (*µ* = 550.0*, σ* = 10.0) and *θ*_2_ ∼ *N* (*µ* = 500.0*, σ* = 10.0)). The black lines represent the posterior distribution. In all plots the *x* axis is time (Ma) whereas the *y* axis is density.

#### 3.2.2 The effect of priors

The bias of endpoint parameters in maximum likelihood estimation is not seen when using Bayesian inference. By applying priors to the *θ.*parameters, the posterior surface can removes the bias in estimation seen in maximum likelihood estimation (Figure 7).

Because of this important role of the priors, the behaviour of the Bayesian model is explored under two situations: Priors for *θ*_1_*, θ*_2_ with relative low and high variance, and priors which are consistent or inconsistent with the true parameter value, that is, whether the true value is near the distribution mean, or very off from it. This allows to verify how sensitive to the prior it is as a function of sample size. When applying priors that are centred on the true parameter values, both low-and high-variance priors are capable of estimating the posterior distributions that include the true value. The effect of the prior variance on the posterior variance however reduces as sample size grows (Figures 7,9).

If an incorrect prior is chosen, one could potentially run into issues where the posterior do not include the true value even for large sample sizes. By applying a prior which is incorrect in the sense that it is centred away from the true values, it is possible to assess the effect of informative and poorly-informative priors on the posterior distriutions. It was found that a low-variance prior which are centred away from the true values have difficulties in converging to the true values even for large sample sizes. This effect is however not seen in high-variance priors, which quickly are capable of converging to the region around the true value as sample size grows (Figures 8,9,10, see panels B, D, F, H for incorrect priors). Overall the method is well behaved as regardless of whether priors are correct or incorrect, informative or poorly-informative, the mean squared error decreases with an increase in sample size (Figure 10). The interquartile ranges tend to increase with sample size only for priors which are incorrect and informative, whereas they always decrease for correct informative and poorly-informative priors, as well as for incorrect poorly-informative priors (Figure 9A,C,E–H).

## 4 Empirical examples

### 4.1 Empirical example 1: The palynomorphs of the Cerrejón Formation

The Cerrejón Formation records the sedimentary dynamics during the Paleocene in northern South America. The nearly 1000 m stratigraphic section is rich in plant macro and microfossils, preserved in a coastal forest environment. Also, it is very important in economic terms, because it preserves multiple coal seams of very high quality, which has been extensively exploited in the Cerrejón open-pit coal mine. Jaramillo et al. (2007) studied the palynomorphs of a long section from the Cerrejón coal mine spanning the Manantial, Cerrejón, Tabaco, and Palmito Formations from base to top. About 195 samples are available in the combined 4752 and 4774 sections, which were identified, counted, and subject to palaeoecological analyses. The data on the 216 different palynomorphs found were herein analysed to illustrate the application of stratigraphic interval estimation, as well as to estimate a distribution for a co-occurrence of stratigraphic intervals. Note that in biostratigraphy, time in the stratigraphic sections (frequently cores) is measured in depth from either top or base. Accordingly, time in this application will represent position in the stratigraphic column instead of absolute time.

#### 4.1.1 Stratigraphic intervals of assorted palynomorphs

Three palynomorphs were chosen for estimating the stratigraphic interval parameters using depth occurrences. Most of the 216 taxa were discarded because they (1) spanned the whole sequence, or (2) had very few occurrences. From the selected palynomorphs, three were chosen which range from 26 to 46 occurrences, and varying in the presence of occurrences close to the top. In two of them, both *θ*_1_*, θ*2 are co-estimated, whereas in a third one *θ*_2_ was fixed instead of co-estimating both endpoint parameters, because the latest occurrences were very close to the top of the stratigraphic column.

The priors on *θ*_1_ ∼ *N* (*µ* = max(occurrences)*, σ* = 140.0), *θ*_2_ ∼ *Exp*(*θ* = min(occurrences)*/*2), and *λ* ∼ *N* (*µ* = 0.0*, σ* = 2.0) were used for each stratigraphic interval. These priors are rather poorly-informative, following the results herein on the possible sensitivity to the prior in the presence of small sample sizes. The parameter *θ*_2_ was fixed to 0.0 in *Ischyosporites problematicus* because the latest occurrences were very close to the top of the section, thus estimating only *θ*1 and *λ*.

The posterior predictive distribution was not calculated in this setting, only the posterior parameter distribution. MCMC sampling was carried out with the NUTS, with 10000 iterations. All parameter traces attained effective sample sizes above 600 (Table 1).

**Table 1:**
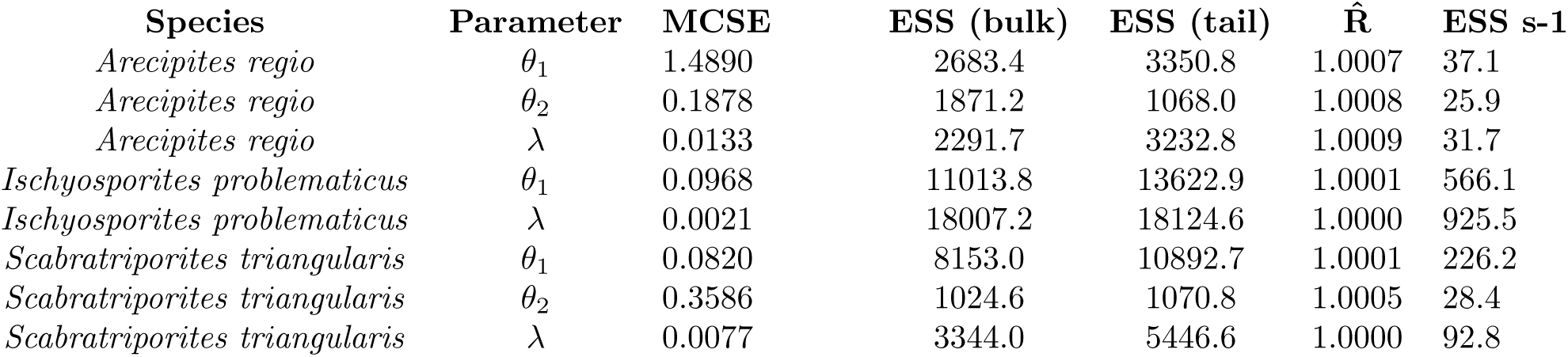
MCMC efficiency and convergence measures for the analyses of the Cerrejón Formation palynomorphs. Abbreviations are Monte-Carlo standard error (MCSE), effective sample size (ESS), effective sample size per second (ESS s-1). 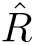 following Vehtari et al. (2021).

Posterior sampling showed good efficiency and convergence (Table 1), and recovered behaviours also seen in simulations. First, stratigraphic intervals where *λ* created a tail towards *θ*_1_ by decreasing the number of occurrences towards the present also made the inference of this parameter to show higher variance than in cases where *λ* was closer to 0.0 (compare Figure 12A and 12B). All posterior distributions of *θ*_1_ were mostly contained into the measured sections (Figure 12). Posterior distribution of *θ*_2_ was bound between 0.0 and the latest occurrence, however, it did not tend to follow the shape of the prior distribution for *Arecipites regio*, thus suggesting a stronger influence of more data (46 occurrences, Table 2, Figure 12A). In contrast, the posterior was similar to the prior in *Scabratriporites triangularis*, presumably due to the smaller sample size (26 occurrences, Table 2, Figure 12C). The posterior distribution of *θ*_2_ was consistently different from the prior for all species, regardless of sample size. The posterior distribution of *λ* were always quite different from 0.0, which means that the stratigraphic interval model is different from uniform.

**Figure 12:**
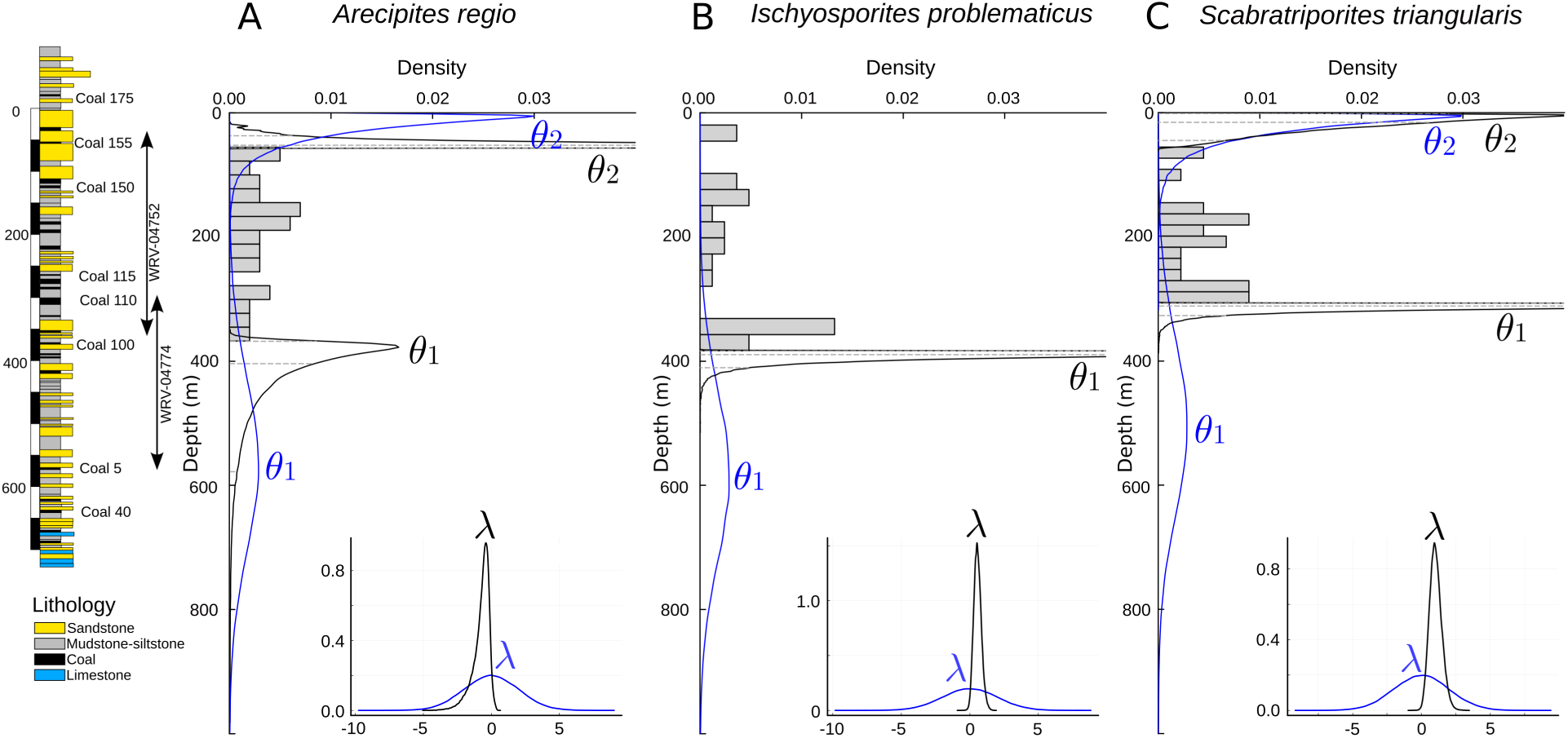
Stratigraphic intervals of A) *Arecipites regio* (*n* = 46), B) *Ischyosporites problematicus* (*n* = 33), and C) *Scabratriporites triangularis* (*n* = 26). The parameter *θ*_2_ was fixed for *Ischyosporites problematicus* as its latest occurrences were very close to the uppermost samples in the stratigraphic column. Offset left is the generalised stratigraphic column of the Cerrejón Formation starting from the top of core WVR-0475 and finishing below core WVR-4774. Subplots A–C are aligned with the generalised column on the depth (m) axis. Lowermost densities correspond to *θ*_1_, uppermost ones to *θ*_2_, and inlay plots to *λ*. Prior distributions are blue, whereas posterior distributions are black. Data are the grey bars in the histogram. Dashed grey lines are the posterior quantiles 0.025, 0.5, and 0.975.

**Table 2:**
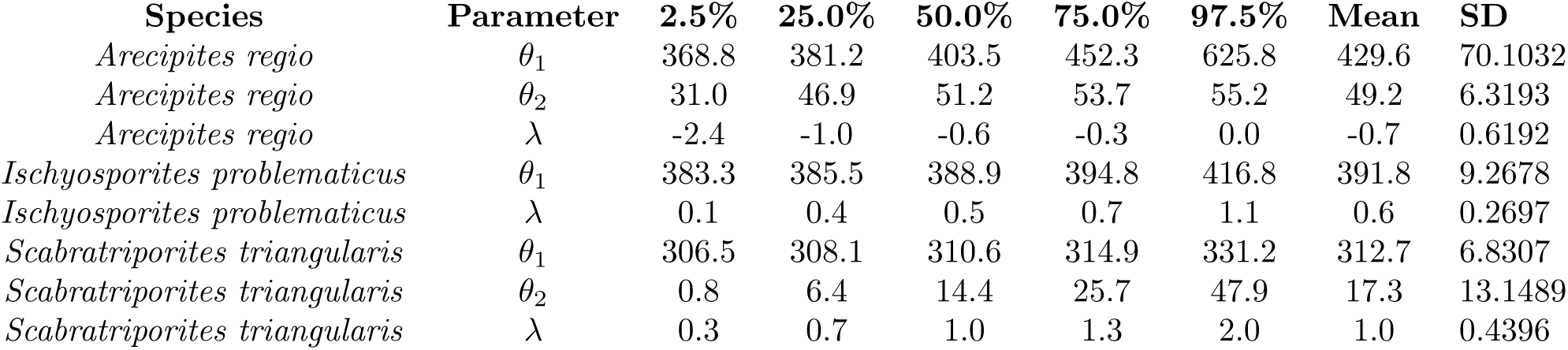
Summary statistics for the posterior sampling of three palynomorphs from the Cerrejón Formation. Percentage columns are quantiles at that given probability value. Abbreviations are standard deviation (SD).

#### 4.1.2 The lost sample of Coal Seam 115

Imagine a fictitious situation: While processing samples collected at the Cerrejón coal mine in northern Colombia, an analyst found out that one important sample had missing information for depth and stratigraphic section in the label, with “Coal Seam 1-something” as the only information available. First-hand stratigraphic information for the sample is unavailable, and therefore the goal is to figure out where in the generalised column the sample came from, as important fossil leaves were also found just at the top of that stratigraphic level and therefore a finer provenance information is key. What is the best estimate of stratigraphic position possible?

After processing the samples for palynomorphs, six of these were found in Coal Seam 115 (true depth 258.99 m which is not known in this fictitious scenario): *Psilatricolporites* sp. (*n* = 70), *Retitricolporites* sp. (*n* = 61), *Psilamonocolpites grandis* (*n* = 44), *Echinatisporis minutus* (*n* = 41), *Retitricolpites* sp. (*n* = 23), and *Matonisporites* sp (*n* = 23). A subset of only occurrences for these six palynomorphs was selected, and the occurrence corresponding to the level 258.88 was removed in order to mimic the absence of information for the position of that level, which reduced in 1 the number of occurrences. The goal here is to first build an age model for each palynomorph using the posterior predictive using the preserved occurrences and then combine these six distributions.

The priors were set as in the previous section. The parameter *θ*_2_ was again fixed to 0.0 in *Psilatricolporites* sp., *Retitricolporites* sp., and *Matonisporites* sp., for the reasons discussed above.

The posterior predictive distribution was calculated here in order to conflate such distribution for each stratigraphic interval into a combined distribution for Coal Seam 115. MCMC sampling was carried out again with the NUTS, but this time using 100000 iterations because 10000 did show issues of convergence for three out of the six species. Afterwards, parameter traces attained effective sample sizes above 600 again. Quantiles were calculated with the area under the curve of the conflated PDF and then compared with the true position for Coal Seam 115.

The use of six palynomorphs for building a conflated distribution for their co-occurrence restricts the possible depth of the mystery sample to the 95% interval 341.8m–54.4m, with a width of 287.4 m. The median of the conflated distribution was 193.8 m. This interval includes the true depth of the Coal seam 115 which is 285.99 m (Figure 13). Four coal seams are found in this interval, 110, 115, 150, and 155. The 95% interval of the conflated distribution was in general restricted to the stratigraphic span of section WRV-4752.

**Figure 13:**
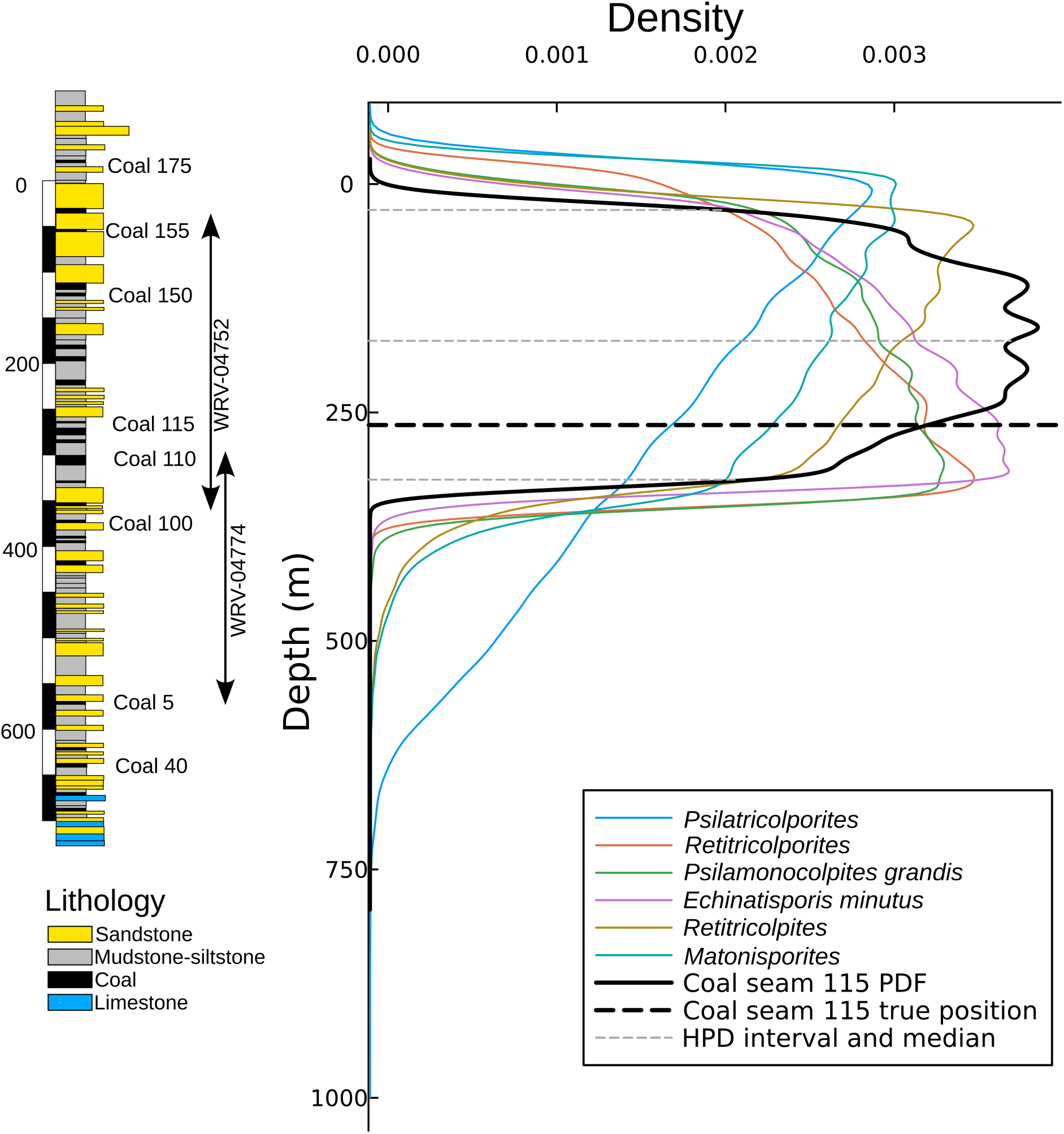
The most probable depth for a mysterious sample labelled Coal Seam 1–something as inferred from the conflation of the posterior predictive distributions of the palynomorphs recovered from that sample. Offset left is the generalised stratigraphic column of the Cerrejón Formation starting from the top of core WVR-0475 and finishing below core WVR-4774. The true depth (Coal Seam 115) falls within the highest density interval of the conflated distribution, although its median is not aligned to the true value.

Most of the palynomorph age models, represented by their posterior predictive distributions, agreed more or less on the interval which determined in great part the shape of the conflation. However, *Psilatricolporites* sp. showed a wider distribution, not contributing much to the final shape because of its greater variance when compared to the posterior predictive distributions of the remaining palynomorphs.

### 4.2 Empirical example 2: The time of origin of barracudas

The origination time of the barracudas (family Sphyraenidae) will be estimated using a curated dataset of fossil occurrences of the family (Ballen, 2020). Because the group is still living, the extinction time *θ*_2_ will be fixed to zero during Bayesian posterior parameter estimation of *θ*_1_ and *λ*. The effect of using different priors was assessed by setting *θ*_1_ ∼ *N* (*µ* = 100.0*, σ* = 25.0), RefOffExp(*s* = 50.0*, o* = 50.0*, ρ* = 1), and *U* (*l* = 50.0*, u* = 300.0). The prior *λ* ∼ *N* (*µ* = 0.0*, σ* = 2.0) was used in all cases. Also, comparisons between removing and leaving duplicate time occurrences were carried out. It is important to note that these duplicates are occurrences from different localities, not redundancies form the same locality, and therefore have the same age just because of uncertainty in dating, and thus are not true duplicates. The effect of this remotion is to reduce sample size.

The posterior predictive distribution was not calculated in this setting, only the posterior parameter distribution. MCMC sampling was carried out with the NUTS, with 10000 iterations. All parameter traces attained effective sample sizes above 600 (Table 3). Simulations were carried out in parallel for efficiency using GNU parallel (Tange, 2011, 2024).

**Table 3:** MCMC sampling efficiency measures for the analyses of the origination time of barracudas. Abbreviations are Monte-Carlo standard error (MCSE), effective sample size (ESS), effective sample size per second (ESS s-1). 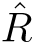 following Vehtari et al. (2021). Clean represents whether duplicates were removed or not.

| Prior | Clean | Parameter | MCSE | ESS (bulk) | ESS (tail) | $\hat{R}$ | ESS s-1 |
| --- | --- | --- | --- | --- | --- | --- | --- |
| N(100.0, 25.0) | No | $\theta_1$ | 0.0828 | 816.7 | 817.8 | 1.0000 | 80.1 |
| N(100.0, 25.0) | No | $\lambda$ | 0.0047 | 1089.7 | 1078.0 | 1.0002 | 106.8 |
| RefOffExp(50.0, 50.0, 1) | No | $\theta_1$ | 0.0346 | 1250.1 | 2268.2 | 1.0000 | 61.5 |
| RefOffExp(50.0, 50.0, 1) | No | $\lambda$ | 0.0028 | 1539.6 | 2141.4 | 1.0009 | 75.7 |
| U(550.0, 300.0) | No | $\theta_1$ | 0.0419 | 1095.2 | 1362.9 | 1.0000 | 53.9 |
| U(550.0, 300.0) | No | $\lambda$ | 0.0032 | 1413.5 | 1910.8 | 1.0003 | 69.5 |
| N(100.0, 25.0) | Yes | $\theta_1$ | 0.4962 | 1287.4 | 889.1 | 1.0001 | 128.8 |
| N(100.0, 25.0) | Yes | $\lambda$ | 0.0252 | 1246.6 | 785.7 | 1.0002 | 124.8 |
| RefOffExp(50.0, 50.0, 1) | Yes | $\theta_1$ | 0.4003 | 1702.3 | 1661.0 | 1.0026 | 95.1 |
| RefOffExp(50.0, 50.0, 1) | Yes | $\lambda$ | 0.0212 | 1599.1 | 1768.0 | 1.0031 | 89.3 |
| U(550.0, 300.0) | Yes | $\theta_1$ | 0.7271 | 1297.7 | 1806.3 | 1.0003 | 73.8 |
| U(550.0, 300.0) | Yes | $\lambda$ | 0.0322 | 1370.2 | 1548.8 | 1.0000 | 77.9 |

Leaving the original dataset with time duplicates (i.e., occurrences with the same time but from different localities, therefore not complete duplicates) results in 75 occurrences. In this scenario the shape of the posterior distribution is quite insensitive to the choice of the prior. Removing duplicates leaves 33 occurrences, for which the choice of the prior impacts somewhat the shape of the posterior distribution.

The most important factors influencing the sensitivity of the posterior seem to be both the prior mean and variance (Fig. 14). The Normal prior tends to have higher density around mean (*σ*^2^ = 625.0), and therefore tends to push more the posterior towards its mean, in this case, into the past. In contrast, the priors RefOffExponential and Uniform, with more variance (*σ*^2^ = 2500.0 and *σ*^2^ = 5208.33 respectively), tend to have a slighter impact on the posterior distribution.

**Figure 14:**
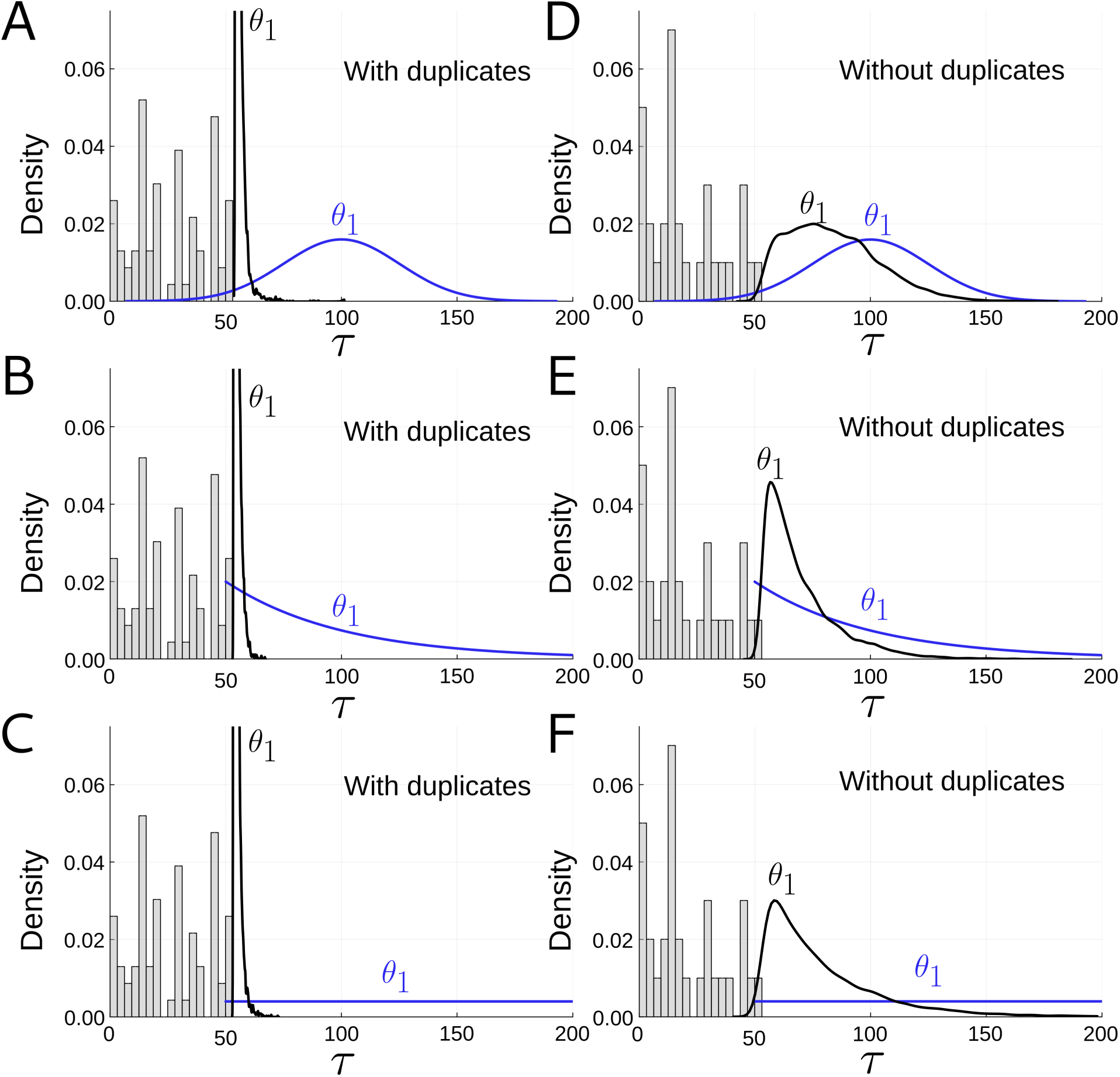
Origination time of the barracudas. Left column are datasets without removing time duplicates. Right column are datasets removing time duplicates. Top row is a prior *θ*_1_ ∼ *N* (100.0, 25.0), middle row is a prior *θ*_1_ ∼ RefOffExp(50.0, 50.0, 1), and bottom row is a prior *θ*_1_ ∼ *U* (50.0, 300.0). Axes represent time (in Ma) on the horizontal, and density on the vertical. Prior distribution in blue, posterior distribution in black. Data are the grey bars in the histogram.

The posterior distribution of *θ*_1_, the origination time, has a highest posterior density interval between 54.3–53.1 Ma (mean age 54.2–54.4 Ma depending on the prior) when using the complete dataset, regardless of the prior used (Table 4). Such interval grows to 113.8–53.8 Ma (narrowest, RefOffExponential prior) and even to 146.4–54.1 Ma (widest, Uniform prior) when using the reduced dataset, this time showing more influence of the prior. In this case, the posterior mean was 70.1 Ma (RefOffExponential prior), 79.1 Ma (Uniform prior), or 83.5 Ma (Normal prior).

**Table 4:** Summary statistics for the posterior sampling of the origination time of barracudas. Percentage columns are quantiles at that given probability value. Clean represents whether duplicates were removed or not. Abbreviations are standard deviation (SD).

| Prior | Clean | Parameter | 2.5% | 25% | 50% | 75% | 97.5% | Mean | SD |
| --- | --- | --- | --- | --- | --- | --- | --- | --- | --- |
| N(100.0, 25.0) | No | $\theta_1$ | 53.1 | 53.4 | 53.8 | 54.8 | 58.8 | 54.4 | 2.0286 |
| N(100.0, 25.0) | No | $\lambda$ | -0.4 | -0.1 | -0.1 | 0.0 | 0.1 | -0.1 | 0.1348 |
| RefOffExp(50.0, 50.0, 1) | No | $\theta_1$ | 53.1 | 53.4 | 53.8 | 54.5 | 57.5 | 54.2 | 1.2295 |
| RefOffExp(50.0, 50.0, 1) | No | $\lambda$ | -0.3 | -0.1 | 0.0 | 0.0 | 0.1 | -0.1 | 0.1090 |
| U(550.0, 300.0) | No | $\theta_1$ | 53.1 | 53.4 | 53.8 | 54.6 | 57.7 | 54.2 | 1.3448 |
| U(550.0, 300.0) | No | $\lambda$ | -0.3 | -0.1 | 0.0 | 0.0 | 0.1 | -0.1 | 0.1133 |
| N(100.0, 25.0) | Yes | $\theta_1$ | 55.3 | 68.3 | 81.1 | 95.8 | 126.4 | 83.5 | 19.2489 |
| N(100.0, 25.0) | Yes | $\lambda$ | -3.9 | -2.5 | -1.7 | -1.1 | -0.3 | -1.8 | 0.9564 |
| RefOffExp(50.0, 50.0, 1) | Yes | $\theta_1$ | 53.8 | 58.5 | 64.9 | 76.3 | 113.8 | 70.1 | 16.3802 |
| RefOffExp(50.0, 50.0, 1) | Yes | $\lambda$ | -3.4 | -1.6 | -1.0 | -0.6 | -0.1 | -1.2 | 0.8516 |
| U(550.0, 300.0) | Yes | $\theta_1$ | 54.1 | 60.9 | 71.3 | 89.3 | 146.4 | 79.1 | 25.0567 |
| U(550.0, 300.0) | Yes | $\lambda$ | -4.4 | -2.2 | -1.3 | -0.8 | -0.2 | -1.6 | 1.1182 |

The posterior distribution of *λ* included zero but showed a prevalence of values on the negative side of the axis (e.g.,-0.4–0.1) for the complete dataset, and was consistently restricted to the negative side of the axis for the reduced dataset (e.g.,-3.4–-1.2). This means that the distribution of occurrences tended to have a tail towards the past, which makes the posterior of *θ*_1_ wider than if it was uniform (cf. Figure 11).

## 5 Discussion

This is one of the first Bayesian methods for stratigraphic intervals to be available in code, and the first Julia package dedicated to estimation of stratigraphic intervals. It is the first to allow flexible specification of a large number of prior distributions, both via the Distributions.jl package or implementing custom ones. Also, the possibility of specifying custom MCMC samplers helps attain convergence very easily. The choice of the NUTS has proven to be quite useful in general cases, and more efficient than the Hamiltonian Monte Carlo, which requires to set more arguments and may run into issues of convergence. The Turing.jl package also provides other samplers including Random-walk Metropolis-Hastings, Gibbs, and compositional Gibbs, thus extending the possibilities of fine-grain control over MCMC sampling. Finally, it allows to fix parameters to given values allows to use stratigraphic intervals in situations where e.g. the lineages in question are extant, but not restricting the user to this as happens with the Adaptive Beta of Wang et al. (2016).

### Maximum likelihood and Bayesian estimation

The bias in estimation where the FAD and LAD are the maximum likelihood estimates is an issue present in other methods for stratigraphic intervals, not just the present one (Strauss and Sadler, 1989; Marshall, 1990; Rivadeneira et al., 2009; Wang et al., 2016, T. Quental, pers. comm.). This makes the use of Bayesian methods more convenient for avoiding biased estimates of origination and extinction times. Wang (2005) studied the maximum likelihood properties of the Four-Parameter Beta distribution and found similar issues with parameter estimation, also advocating for constraining the likelihood surface using poorly-informative priors on the endpoint parameters *θ*_1_*, θ*_2_. He discussed the conditions under which parameter estimation is possible, through the use of Bayesian methods. The approach herein is in line with his findings, but is of more general application as it does not enforce the use of a specific prior on endpoint parameters. Wang (2005) examined the correlation between parameters and concluded that endpoints tend to be correlated. This is an issue for inference using Metropolis-Hasting samplers, and it was unnecessary to examine it here because both the Hamiltonian and NUTS are very efficient traversing surfaces in these situations.

### Priors

The behaviour of different priors was in general terms similar in that given sufficient data, the likelihood function is capable of dominating the information content, thus determining to a great extent the parameter posterior distributions. However, it is clear that even fairly poorly-informative priors can exert some influence when sample size is small (Figure 14). It is important to note that priors which are very off form the true values (herein called incorrect priors) have an undesirable influence on the posterior distributions, which is proportional to the information content in the prior distribution, and therefore inversely proportional to the variance. Thus, even if a prior is incorrect because its mean is off from the true value, it is possible to make it behave more properly by setting a larger variance for it, that is, making it fairly poorly-informative. These findings suggest that the method is robust to prior specification, and it is recommended that empirical applications with small sample size (around an order to 50 occurrences or less) specify wide priors in order to avoid any undesirable influence of the prior on the posterior. Although priors which are both incorrect and informative behave very badly in the sense that they do not recover the true value in simulations, they tend to approach the true values which may eventually converge if letting simulations to run for very long and with very large sample sizes (above 500 in this study).

### Comparison with other methods

The Three-Parameter Beta model, in the Bayesian formulation, shares features with several methods already available. On one side, it assumes continuous sampling, as is common to available tools, but in contrast with methods for discrete sampling (Weiss and Marshall, 1999; Weiss et al., 2004). It can accommodate both uniform (Strauss and Sadler, 1989; Solow, 1993a; Ferraz et al., 2003; Weiss and Marshall, 1999; McInerny et al., 2006; Wang et al., 2016) and non-uniform (Solow, 1993b; Marshall, 1997; McCarthy, 1998; Weiss et al., 2004; Ferraz et al., 2003; Wang et al., 2016) preservation potentials, thus being similar to many Class 1 and Class 2 methods, and in contrast with non-parametric or distribution free methods (Marshall, 1994; Solow and Roberts, 2003). The method can condition on a fixed endpoint parameter like many do (e.g., Wang et al., 2016), in particular those tailored for sighting data in conservation biology, which assume that the oldest occurrence is an arbitrary zero point, and then estimate the extinction time (Ferraz et al., 2003; Solow, 2005; Kodikara et al., 2020), but it is not forced to such conditioning and therefore can estimate both the origination and extinction times like some methods from palaeobiology (Strauss and Sadler, 1989; Marshall, 1994). It is unique however in allowing the user to specify any prior they want, which is supported by the Distributions.jl package or implemented in StratIntervals.jl (e.g., the RefOffExponential distribution). Also, it is unique among many Bayesian methods in that it is fully implemented in a software package which can be easily integrated with others from the Julia language ecosystem. This helps in allowing the user to extend the models available into three directions: Choice of priors, which multiple works have found to matter (Alroy, 2016a,b; Solow, 2016a,b), choice of samplers, therefore allowing to improve sampling efficiency, and the possibility to extend methods by using different likelihood functions.

### Empirical applications

The combined effect of the tail of the distribution towards the past, as well as the lower sample size in the reduced dataset explains why the posterior distribution of *θ*_1_ was considerably wider, and should be considered when assessing the results of the application of the present method to empirical datasets. Although the choice of the prior for *θ*_1_ has some influence on the posterior distribution, this is greatly reduced by sample size, under the same inferential settings. If the distribution of occurrences is concentrate towards any of the endpoints, the posterior estimation of the opposite endpoint will suffer from wider variance. This effect is seen both in the palynomorph and the barracudas datasets. This is a feature of the data and the distribution rather than a pathological behaviour of the method. It is comparable to trying to fit a curve to an asymmetric distribution, variations on the tailed side can vary to a larger extent than the part attached to the other side. This makes sense as more of the density of the information in the data are concentrated to the other side of the distribution, and therefore having less information on the tailed side (Figures 12 and 14).

The conflation of distributions showed desirable properties in real-world applications. The fact that a reasonable position for the Coal seam 115 was recovered just by observing the occurrences of only six palynomorphs present in the sample is exciting, as normally much more taxa are available in real-world samples for biostratigraphic analyses (Jaramillo et al., 2007). Its potential however is not restricted to micropalaeontology but instead to any application where combinations of distributions is of interest. For instance, Ballen and Reinales (2025) showed its application to the construction of secondary calibrations in divergence time estimation. Although methods for stratigraphic intervals were born from the field of quantitative palaeobiology, their application to questions in evolutionary biology and related fields can be much wider, especially in the field of divergence time estimation.

Calculating the co-occurrence of multiple lineages in time has shown to be useful in estimating the time of co-occurrence of collections of objects in time. This method does not require the intervals to represent living organisms, and therefore can be easily applied to historical objects such as musical pieces which are preserved in multiples sources, e.g. Gregorian chant melodies in manuscripts (Ballen et al., 2026). These scenarios are quite common in the study of cultural evolution and digital humanities, which are interesting fields where data sometimes correspond to series of occurrences in time, the methods presented herein are promising.

## 6 Conclusions

1. The Three-parameter Beta distribution has been shown to be useful for estimating endpoints in stratigraphic intervals. However, its likelihood function presents the same issues of a larger family of nonregular models where endpoint parameters define the support of the function. This issue however disappears when using a Bayesian formulation of the model
2. The Bayesian method has been shown to be useful in estimating stratigraphic intervals themselves, but also extending these to continuous representations of a lineage in time, and also when estimating the co-occurrence of such lineages.
3. The analytical framework herein proposed is expected to have a wider impact both in palaeobiology and evolutionary biology by providing tools for studying lineages through time without the need of explicit phylogenies and taking the most out of imperfect fossil record.

## Acknowledgments

This research was supported by FAPESP through a doctoral scholarship (2014/11558-5), an internship abroad (2016/02253-1), and a postdoctoral fellowship (2023/07838-1), as well as by the BBSRC (grant BB/T01282X/1 awarded to Mario dos Reis). Thanks to the editor and reviewers for all their helpful feedback. I thank Mario dos Reis and Ziheng Yang for valuable feedback during early phases of this project, I could not have finished it without their guidance and criticism in methods development. I thank Phil Donoghue, Davide Pisani and the Molecular Palaeobiology Group at University of Bristol for inviting me to deliver a talk on the topic of this paper when it was still in early phases of development. I thank Tiago Quental and Gustavo Burin for discussions on methods for palaeobiology and macroevolution throughout the years, and for early discussions on methods for stratigraphic intervals since my first year of doctoral studies. I thank Carlos Jaramillo for supporting my career in palaeontology, and for encouraging me and several other Colombian students to build analytical and computational skills in palaeobiology. Discussions on palaeontology and geology with many colleagues (Javier Luque, Jorge W. Moreno, Camila Martinez, Maria Camila Vallejo, Federico Moreno, Andŕes Ćardenas, Edwin Cadena, Camilo Montes, Catalina Suarez, and Sandra Reinales) have been a valuable source of inspiration. Special thanks to Henrique Rieger for thoroughly reading the manuscript, catching errors, and providing suggestions to improve it. Living ammonite reconstruction in Figure 1 thanks to Kouhei Futaka, available under Creative Commons license (https://doi.org/10.7875/togopic.2020.08), and ammonite drawing in Figure 1 thanks to Izolende, also available under Creative Commons (https://commons.wikimedia.org/wiki/User:Ilzolende). Special thanks to Jorge W. Moreno-Bernal for the ammonite reconstructions in Figure 5. Finally, I thank Sandra Reinales for supporting me personally and professionally in this and many other projects throughout the years.

## Author’s contribution

The author contributed to conceptualisation, development, and writing.

## Code and data statement

All the simulation code is available at https://github.com/gaballench/bayesian_collection_stratintervals, as well as on zenodo (DOI: 10.5281/zenodo.14868381).

## References

Alroy, J. (2014). A simple Bayesian method of inferring extinction. Paleobiology, 40(4):584–607.

Alroy, J. (2015). Current extinction rates of reptiles and amphibians. Proceedings of the National Academy of Sciences, 112(42):13003–13008.

Alroy, J. (2016a). A simple Bayesian method of inferring extinction: Reply. Ecology, 97(3).

Alroy, J. (2016b). Reply to Solow: Sense and nonsense in the choice of extinction priors. Proceedings of the National Academy of Sciences, 113(9):E1133–E1133.

Ballen, G. A. (2020). New records of the genus *Sphyraena* (Teleostei: Sphyraenidae) from the Caribbean with comments on dental characters in the genus. Journal of Vertebrate Paleontology, 40(6):e1849246.

Ballen, G. A., Mühlová, K. H., and Hajič Jr, J. (2026). What did the dove sing to pope gregory? ancestral melody reconstruction in gregorian chant using bayesian phylogenetics. PloS one, 21(5):e0350014.

Ballen, G. A. and Reinales, S. (2025). tbea: tools for pre-and post-processing in bayesian evolutionary analyses. Evolutionary Journal of the Linnean Society, 4(1):kzaf017.

Besançon, M., Papamarkou, T., Anthoff, D., Arslan, A., Byrne, S., Lin, D., and Pearson, J. (2021). Distributions.jl: Definition and modeling of probability distributions in the JuliaStats Ecosystem. Journal of Statistical Software, 98(16):1–30.

Bezanson, J., Edelman, A., Karpinski, S., and Shah, V. B. (2017). Julia: A fresh approach to numerical computing. SIAM Review, 59(1):65–98.

Boakes, E. H., Rout, T. M., and Collen, B. (2015). Inferring species extinction: The use of sighting records. Methods in Ecology and Evolution, 6(6):678–687.

Bouchet-Valat, M. and Kamiński, B. (2023). DataFrames.jl: Flexible and fast tabular data in Julia. Journal of Statistical Software, 107(4):1–32.

Delignette-Muller, M. L. and Dutang, C. (2015). fitdistrplus: An r package for fitting distributions. Journal of statistical software, 64:1–34.

Ebrahimi, N., Maasoumi, E., and Soofi, E. S. (1999). Ordering univariate distributions by entropy and variance. Journal of Econometrics, 90(2):317–336.

Ferraz, G., Russell, G. J., Stouffer, P. C., Bierregaard Jr, R. O., Pimm, S. L., and Lovejoy, T. E. (2003). Rates of species loss from Amazonian forest fragments. Proceedings of the National Academy of Sciences, 100(24):14069–14073.

Ge, H., Xu, K., and Ghahramani, Z. (2018). Turing: A language for flexible probabilistic inference. In International Conference on Artificial Intelligence and Statistics, AISTATS 2018, 9-11 April 2018, Playa Blanca, Lanzarote, Canary Islands, Spain, pages 1682–1690.

Genest, C. and Zidek, J. V. (1986). Combining probability distributions: A critique and an annotated bibliography. Statistical Science, 1(1):114–135.

Hall, P. and Wang, J. Z. (2005). Bayesian likelihood methods for estimating the end point of a distribution. Journal of the Royal Statistical Society Series B: Statistical Methodology, 67(5):717–729.

Hill, T. P. (2011). Conflations of probability distributions. Transactions of the American Mathematical Society, 363(6):3351–3372.

Hill, T. P. and Miller, J. (2011). How to combine independent data sets for the same quantity. Chaos: An Interdisciplinary Journal of Nonlinear Science, 21(3):033102.

Hoffman, M. D. and Gelman, A. (2014). The No-U-Turn sampler: Adaptively setting path lengths in Hamiltonian Monte Carlo. Journal of Machine Learning Research, 15(1):1593–1623.

Holland, S. M., Patzkowsky, M. E., and Loughney, K. M. (2024). Stratigraphic Paleobiology. Paleobiology, page 1–18.

Jaramillo, C. A., Bayona, G., Pardo-Trujillo, A., Rueda, M., Torres, V., Harrington, G. J., and Mora, G. (2007). The palynology of the Cerrejón Formation (upper Paleocene) of northern Colombia. Palynology, 31(1):153–189.

Johnson, S. G. (2013). QuadGK.jl: Gauss–Kronrod integration in Julia. https://github.com/JuliaMath/ QuadGK.jl.

Kodikara, S., Demirhan, H., Wang, Y., Solow, A., and Stone, L. (2020). Inferring extinction year using a Bayesian approach. Methods in Ecology and Evolution, 11(8):964–973.

Lehmann, E. L. and Casella, G. (1998). Theory of point estimation, Second Edition. Springer.

Lindley, D. V. (1972). Bayesian statistics: A review. SIAM.

Marshall, C. R. (1990). Confidence intervals on stratigraphic ranges. Paleobiology, 16(1):1–10.

Marshall, C. R. (1994). Confidence intervals on stratigraphic ranges: Partial relaxation of the assumption of randomly distributed fossil horizons. Paleobiology, 20(4):459–469.

Marshall, C. R. (1997). Confidence intervals on stratigraphic ranges with nonrandom distributions of fossil horizons. Paleobiology, 23(2):165–173.

Marshall, C. R. (2010). Using confidence intervals to quantify the uncertainty in the end-points of stratigraphic ranges. Quantitative Methods in Paleobiology, 16:291–316.

McCarthy, M. A. (1998). Identifying declining and threatened species with museum data. Biological conservation, 83(1):9–17.

McCrea, R. S., Cheale, T., Campillo-Funollet, E., and Roberts, D. L. (2024). Inferring species extinction from sighting data. Cambridge Prisms: Extinction, 2:e19.

McInerny, G. J., Roberts, D. L., Davy, A. J., and Cribb, P. J. (2006). Significance of sighting rate in inferring extinction and threat. Conservation biology, 20(2):562–567.

Rivadeneira, M. M., Hunt, G., and Roy, K. (2009). The use of sighting records to infer species extinctions: An evaluation of different methods. Ecology, 90(5):1291–1300.

Sareen, S. (2003). Reference bayesian inference in nonregular models. Journal of Econometrics, 113(2):265–288.

Shannon, C. E. (1948). A mathematical theory of communication. The Bell system technical journal, 27(3):379–423.

Signor, P. W. and Lipps, J. H. (1982). Sampling bias, gradual extinction patterns and catastrophes in the fossil record. Geological Society of America Special Paper, 190:291–296.

Silverman, B. W. (1998). Density estimation for statistics and data analysis.

Smith, J., Dowding, E., Abdelhady, A., Abondio, P., Araujo, R., Aze, T., Balisi, M., Buatois, L., Carvajal-Chitty, H., Chattopadhyay, D., Coiro, M., Dietl, G., Gonzalez Arango, C., Kevrekidis, C., Kimmig, J., Mychajliw, A., Pimiento, C., Regalado Fernandez, O., K.M., S., Warnock, R., Yang, T., Yasuhara, M., Akita, L., Allen, B., Anderson, B., Andreoletti, J., Archuby, F., Ballen, G., Bari, M., Benton, M., Bergh, E., Brambilla, L., Brombacher, A., Chan, Y., Chiarenza, A., Chinzorig, T., Coates, K., Cordie, D., Cortes-Sanchez, M., Cruz-Vega, E., Cybulski, J., De Baets, K., De Entrambasaguas, J., Dillon, E., Du, A., Dunhill, A., Erlandson, J., Forel, M., Foster, W., Gates, T., Gavryushkina, A., Grace, M., Grossart, H., Hansel, P., Harnik, P., Hopkins, M., Hopkins, S., Hu, K., Huang, H., Irmis, R., Jaques, V., Jenkins, X., Jukar, A., Kelley, P., Kihn, R., Klompmaker, A., Kocsis, A., Kriwet, J., Lazarus, D., Liao, C., Lin, C., Louys, J., Lozano-Fernandez, J., Lozano-Francisco, M., Lueders-Dumont, J., Malve, M., Martindale, R., Mazzini, I., Modenini, G., Mondal, S., Mondini, M., Monferran, M., Mulvey, L., Nanglu, K., Nguyen, J., Norris, R., O’Dea, A., Ollendorf, A., Orihuela, J., Pandolfi, J., Pereira, T., Piro, A., Plotnick, R., Plaza-Torres, S., Porto, A., Prieto-Marquez, A., Punyasena, S., Quental, T., Raja, N., Ranaivosoa, V., Ribas-Deulofeu, L., Rivals, F., Roden, V., Rosso, A., Saleh, F., Salvador, R., Saupe, E., Schneider, S., Sclafani, J., Smith, M., Souron, A., Steinbauer, M., Stewart, M., Tambussi, C., Thomas, E., Tschopp, E., Tutken, T., Varela, S., Vezzosi, R., Villasenor, A., Weinkauf, M., Zanno, L., Zhang, C., Zhao, Q., and Kiessling, W. (2025). Big questions in paleontology: A community-driven project to motivate new insights about the history of life on Earth. Paleobiology, 51(3):408–431.

Solow, A. R. (1993a). Inferring extinction from sighting data. Ecology, 74(3):962–964.

Solow, A. R. (1993b). Inferring extinction in a declining population. Journal of Mathematical Biology, 32:79–82.

Solow, A. R. (2005). Inferring extinction from a sighting record. Mathematical biosciences, 195(1):47–55.

Solow, A. R. (2016a). A simple Bayesian method of inferring extinction: Comment. Ecology, 97(3).

Solow, A. R. (2016b). On Bayesian inference about extinction. Proceedings of the National Academy of Sciences, 113(9):E1132–E1132.

Solow, A. R. and Roberts, D. L. (2003). A nonparametric test for extinction based on a sighting record. Ecology, 84(5):1329–1332.

Strauss, D. and Sadler, P. M. (1989). Classical confidence intervals and Bayesian probability estimates for ends of local taxon ranges. Mathematical Geology, 21(4):411–427.

Tange, O. (2011). Gnu parallel–the command-line power tool. USENIX Mag, 36(1):42.

Tange, O. (2024). GNU Parallel 20240522 (’Tbilisi’). 10.5281/zenodo.11247979.

Vehtari, A., Gelman, A., Simpson, D., Carpenter, B., and Bürkner, P.-C. (2021). Rank-normalization, folding, and localization: An improved *R*^^^ for assessing convergence of MCMC. Bayesian analysis, 16(2):667–718.

Wang, J. Z. (2005). A Note on Estimation in the Four-Parameter Beta Distribution. Communications in Statistics - Simulation and Computation, 34(3):495–501.

Wang, S. C., Everson, P. J., Zhou, H. J., Park, D., and Chudzicki, D. J. (2016). Adaptive credible intervals on stratigraphic ranges when recovery potential is unknown. Paleobiology, 42(2):240–256.

Weiss, R. E., Basu, S., and Marshall, C. R. (2004). A framework for analysing fossil record data. In Tools for constructing chronologies: crossing disciplinary boundaries, pages 213–230. Springer.

Weiss, R. E. and Marshall, C. R. (1999). The uncertainty in the true end point of a fossil’s stratigraphic range when stratigraphic sections are sampled discretely. Mathematical Geology, 31:435–453.

