## Supplementary for "A flexible Bayesian method for estimating stratigraphic intervals and their co-occurrence in time"

Gustavo A. Ballen[1,2,3,4,*]

[1] Instituto de Biociências de Botucatu, Universidade Estadual Paulista "Júlio de Mesquita Filho", Botucatu, SP, Brazil
[2] School of Biological and Behavioural Sciences, Queen Mary University of London, London, United Kingdom
[3] Center for Tropical Paleoecology and Archaeology, Smithsonian Tropical Research Institute, Ancón, Balboa, Panamá
[4] Museu de Zoologia da Universidade de São Paulo, Universidade de São Paulo, São Paulo, SP, Brazil
Orcid https://orcid.org/0000-0001-5424-8608

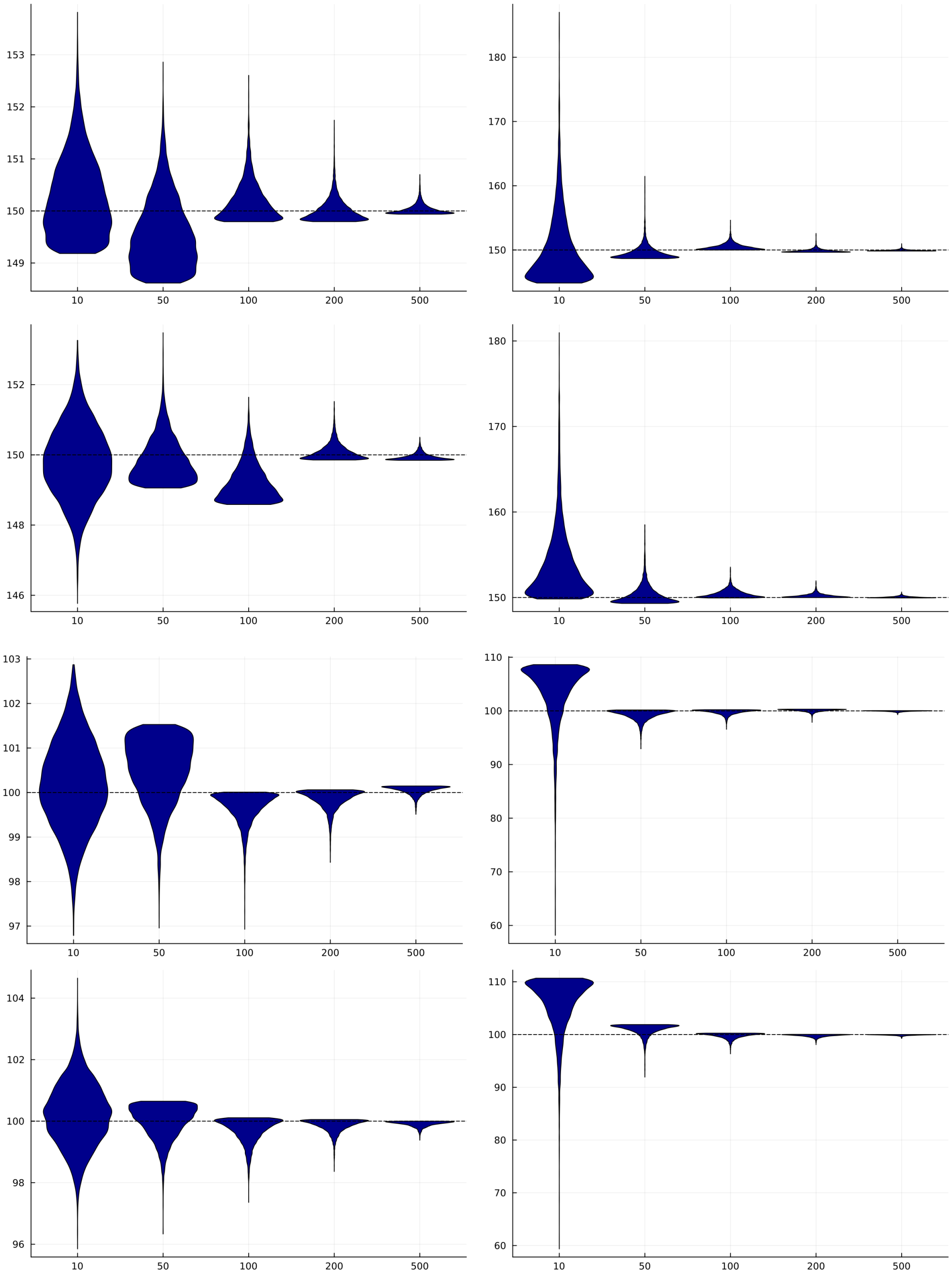


Figure 1: Posterior distribution of a single simulated dataset of varying sample size from the ThreeParBeta$(\tau; \theta_1, \theta_2, \lambda)$ distribution to illustrate the effect of sample size ($x$ axis, in Ma) and prior variance on $\theta_1$ and $\theta_2$. Simulations were carried out by fixing or co-estimating the endpoint parameters. Top left, posterior distribution of $\theta_1$ with prior $N(\mu = 150.0, \sigma = 1.0)$, and then the next one co-estimating both but plotting just $\theta_1$. Middle third left, posterior distribution of $\theta_2$ with prior $N(\mu = 100.0, \sigma = 1.0)$, bottom left the same but co-estimating endpoint parameters and plotting $\theta_2$. Top right, posterior distribution of $\theta_1$ with prior $N(\mu = 150.0, \sigma = 20.0)$ and then the next one co-estimating both but plotting just $\theta_1$. Middle third left, posterior distribution of $\theta_2$ with prior $N(\mu = 100.0, \sigma = 20.0)$, bottom left the same but co-estimating endpoint parameters and plotting $\theta_2$. The dashed line represents the true value of the parameter in all plots.

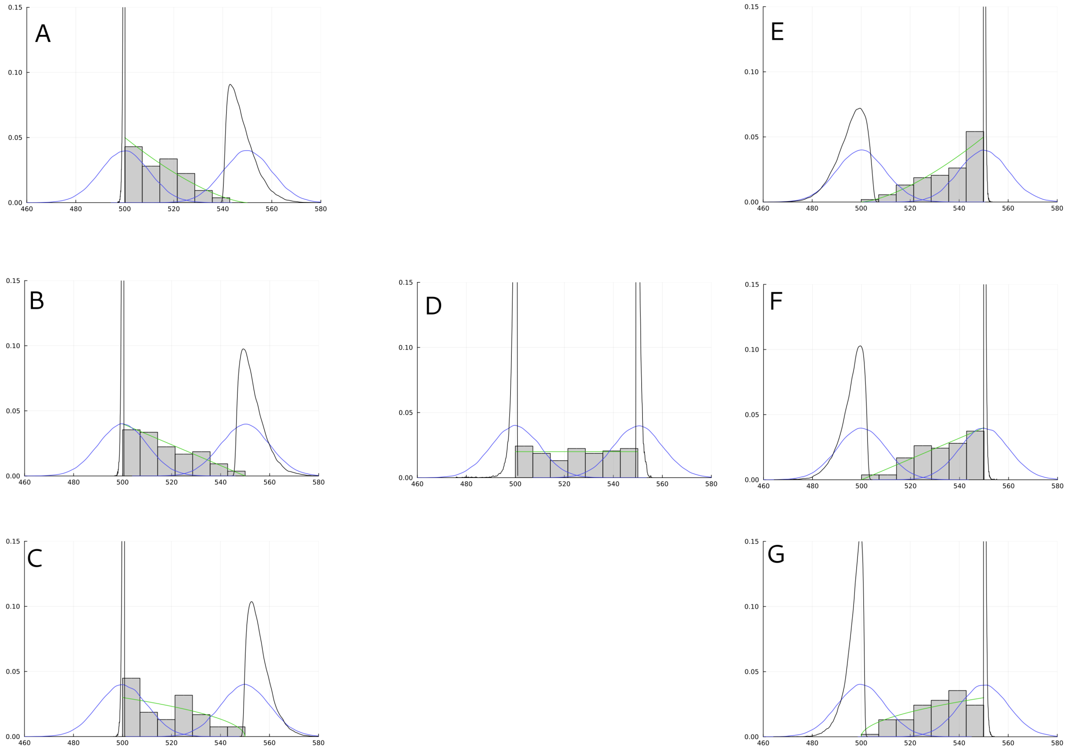


Figure 2: Effect of $\lambda$ on $\theta_1$ and $\theta_2$ for a sample size of 75 occurrences sampled from different parameter values of $\lambda$: A) -1.5, B) -1.0, C) -0.5, D) 0.0, G) 0.5, F) 1.0, E) 1.5. In all plots the green line represents the ThreeParBeta($\tau; \theta_1 = 550.0, \theta_2 = 500.0, \lambda$) from where the 75 occurrences were sampled. Blue lines represent the rather uninformative prior centered at the true value on each ($\theta_1 \sim N(\mu = 550.0, \sigma = 10.0)$ and $\theta_2 \sim N(\mu = 500.0, \sigma = 10.0)$). The black lines represent the posterior distribution. In all plots the $x$ axis is time (Ma) whereas the $y$ axis is density.
